# Optimizing the molecular diagnosis of Covid-19 by combining RT-PCR and a pseudo-convolutional machine learning approach to characterize virus DNA sequences

**DOI:** 10.1101/2020.06.02.129775

**Authors:** Juliana Carneiro Gomes, Aras Ismael Masood, Leandro Honorato de S. Silva, Janderson Ferreira, Agostinho A. F. Júnior, Allana Lais dos Santos Rocha, Letícia Castro, Nathália R. C. da Silva, Bruno J. T. Fernandes, Wellington Pinheiro dos Santos

**Affiliations:** Programa de Engenharia da Computação, Escola Politécnica da Universidade de Pernambuco, POLI-UPE, Recife, Brazil; Information Technology Department, Technical College of Informatics, Sulaimani Polytechnic University, Sulaymaniyah, Iraq; Instituto Federal de Educação, Ciência e Tecnologia da Paraíba, Campus Cajazeiras, IFPB, Cajazeiras, Brazil; Departamento de Engenharia Biomédica, Universidade Federal de Pernambuco, DEBM-UFPE, Recife, Brazil

**Keywords:** Covid-19 diagnosis, Covid-19 molecular diagnosis, DNA sequences representation, pseudo-convolutional methods, RT-PCR optimization

## Abstract

The proliferation of the SARS-Cov-2 virus to the whole world caused more than 250,000 deaths worldwide and over 4 million confirmed cases. The severity of Covid-19, the exponential rate at which the virus proliferates, and the rapid exhaustion of the public health resources are critical factors. The RT-PCR with virus DNA identification is still the benchmark Covid-19 diagnosis method. In this work we propose a new technique for representing DNA sequences: they are divided into smaller sequences with overlap in a pseudo-convolutional approach, and represented by co-occurrence matrices. This technique analyzes the DNA sequences obtained by the RT-PCR method, eliminating sequence alignment. Through the proposed method, it is possible to identify virus sequences from a large database: 347,363 virus DNA sequences from 24 virus families and SARS-Cov-2. Experiments with all 24 virus families and SARS-Cov-2 (multi-class scenario) resulted 0.822222 *±* 0.05613 for sensitivity and 0.99974 *±* 0.00001 for specificity using Random Forests with 100 trees and 30% overlap. When we compared SARS-Cov-2 with similar-symptoms virus families, we got 0.97059 *±* 0.03387 for sensitivity, and 0.99187 *±* 0.00046 for specificity with MLP classifier and 30% overlap. In the real test scenario, in which SARS-Cov-2 is compared to Coronaviridae and healthy human DNA sequences, we got 0.98824 *±* 001198 for sensitivity and 0.99860 *±* 0.00020 for specificity with MLP and 50% overlap. Therefore, the molecular diagnosis of Covid-19 can be optimized by combining RT-PCR and our pseudo-convolutional method to identify SARS-Cov-2 DNA sequences faster with higher specificity and sensitivity.

## 1. Introduction

At the end of 2019, the proliferation of the SARS-Cov-2 virus appeared in the city of Wuhan, China (Zhou et al., 2020). In a few months, there are more than 250,000 deaths worldwide and over 4 million confirmed cases (WHO, 2020b). Covid-19, as it became known, is a respiratory syndrome. In moderate cases, it manifests clinically as pneumonia. In critical cases, a disease can lead to respiratory failure, septic shock, and/or multiple organ dysfunction (MOD) or failure (MOF) (Cascella et al., 2020; Peeri et al., 2020; Wang et al., 2020a).

Besides the severity of the disease, the exponential rate at which the virus proliferates is an aggravating factor. The transmission of the virus often occurs through asymptomatic people. The contagion is given by drops or secretions from sneezing or coughing (Cascella et al., 2020). Because of this, many countries have been experiencing overcrowding in their hospital centers. Most medical professionals are working long hours, and the number of pulmonary ventilators is not enough for all patients. This scenario has led dozens of countries to adopt measures of social isolation. They attempt to contain the dissemination, and to mitigate the number of people who need hospitalization (Hellewell et al., 2020; Wilder-Smith & Freedman, 2020; Kraemer et al., 2020).

In response to this growing pandemic, several companies and research centers worldwide have researched and developed methods for diagnosing Covid-19 (Wang et al., 2020b). Among them, rapid tests emerged, which can provide results in about 30 minutes. One type of rapid test is the Rapid Diagnostic Test (RDT). Through samples from the patient’s respiratory tract, RDT seeks to detect the presence of antigens. Antigens are substances that are foreign to the body, causing immune responses. These responses produce specific antibodies, capable of binding to and interacting with the antigen, ensuring the protection of the organism. Thus, in tests of the RDT type, antibodies are fixed on paper tapes and placed in plastic capsules, similar to the well-known pregnancy tests. If the target antigen is present in the patient’s sample at certain concentrations, it will attach to the antibodies on the tape, generating a visual signal. Unfortunately, this method has some restrictions. First, it is only possible to detect in the acute stages of infection, when antigens are expressed. In addition, efficiency depends on factors such as quality and the collection protocol and the formulation of reagents. We must also emphasize that the possibility of false positives, when the antibodies present on the tape recognize antigens from other types of viruses. For these reasons, the sensitivity of the RDT can vary from 34 to 80% (Bruning et al., 2017; WHO, 2020a).

Another type of rapid test is based on host antibody detection. In this case, antibodies are detected in the patient’s blood samples, depending on factors such as age, nutrition, disease severity and medications. However, recent studies have shown that the immune response is very weak, late or even absent in many cases of patients confirmed with Covid-19 (Döhla et al., 2020; Patel et al., 2020; Burog et al., 2020; Li et al., 2020; Liu et al., 2020; Zhang et al., 2020; Pan et al., 2020). This means that this type of detection is often only possible in cases of recovered patients. The study (Long et al., 2020) reports 285 patients who tested positive for IgG. However, these immune responses were seen 19 days after the first symptoms. This condition makes testing ineffective in many situations, as opportunities for treatment and clinical interventions no longer exist. Therefore, WHO does not currently recommend these types of rapid diagnostic tests for Covid-19. The suggestion is to use them in research contexts or as a way of screening patients, or of potential diagnosis (WHO, 2020a).

Therefore, the benchmark for Covid-19 diagnosis is molecular diagnosis or RT-PCR with DNA sequencing and identification (Patel et al., 2020; Tahamtan & Ardebili, 2020). Throat swab samples are usually collected from suspected patients in this type of analysis. The samples are then placed in tubes with virus preservation solutions, where the genetic material of the virus can be extracted. In this case, the single-stranded RNA. In the first phase, reverse transcription occurs, where a complementary DNA molecule (cDNA) to the virus RNA is synthesized. This process takes place through the DNA polymerase enzyme. The RNA is then removed, and the Taq DNA polymerase enzyme produces double-stranded DNA, which is a copy of the virus’s RNA. Then, the PCR exponentially amplifies fragments of this DNA during successive cycles, generating millions of copies to be analyzed. In the following, the cDNA is aligned with sequences from the SARS-Cov2 virus. Sequence alignment is a traditional method for analyzing similarity between sequences. Among the most consolidated methods are BLAST and FASTA. If there is a match between both sequences, then the patient is confirmed positive. Otherwise, the patient is considered negative for Covid-19 (Bosco & Di Gangi, 2016; Rizzo et al., 2015; Zhang & Harmon, 2020; Chan et al., 2020).

Although so far RT-PCR with DNA identification is considered the most accurate and effective method, there are still some weaknesses. A major limitation of the sequence alignment methods is the computational complexity and time consumption. In many cases, patients can take days to receive the diagnosis due to sample preparation and genomic analysis. Because of this, several studies have proposed alignment free methods for genomic sequences classification. Most of these methodologies involve a feature extraction method such as spectral representation of DNA sequences. Thus, the representative attributes of the sequence can be combined with methods of artificial intelligence, especially machine learning. This makes possible to separate each analyzed sequence into a class (Covid-19 positive or Covid-19 negative, for example) (Bosco & Di Gangi, 2016; Rizzo et al., 2015).

In this work we propose a new technique for representing sequences based on the analysis of the relationships between nitrogenous bases. This technique analyzes the DNA sequences obtained by the RT-PCR method, eliminating the alignment process. The idea is as follows: a DNA sequence is divided into *n* smaller sequences. Each subsequence *i* is superimposed with a part of the subsequence *i−*1 and with a part of the subsequence *i*+1, giving rise to two new subsequences. These smaller sequences are represented by co-occurrence matrices. The matrices are square with 4*x*4 dimensions, with number of rows and columns corresponding to each of the nitrogenous bases of DNA (Adenine, Cytosine, Thymine, and Guanine). The co-occurrence matrix considers the occurrence of each of the bases, as well as the relationship between bases and their immediate neighbors. Then, the co-occurrence matrices are stacked together, forming a volume. Considering that the sequences can be subdivided into smaller and smaller subsets, with the formation of new co-occurrence matrices, the proposed method has a pseudo-convolutional aspect from the algorithmic point of view. After obtaining the set of matrices, they are then concatenated, forming attribute vectors. These extracted attributes correspond to a high-level vectorial representation of the initial DNA sequence, independent from the size of the sequence. This feature vector can be classified by machine learning techniques.

Through the proposed method, it is possible to identify virus sequences from a relatively large database. Several advantages can be pointed out with this approach: First, it is not necessary to pre-align the sequence under investigation in relation to the reference sequences; Second, the sequence under study is compared with a wide set of sequences of given classes, and not just with a reference sequence, strengthening the reliability of the test. We also emphasize that the method can be applied to sequences of any size.

The present work seeks to describe and test the new method of feature extraction to represent sequences of nitrogenous bases. Our main objective is to optimize the RT-PCR, the benchmark for Covid-19 diagnosis. To reach this goal, we used genomic sequences of different viruses obtained in the repository VIPR (Virus Pathogen Resource) Pickett et al. (2012). We used 24 virus families with more than 500 sequences each, including the SARS-Cov2 family. Each sequence was submitted to the representation process described here. In the following, we performed multiple experiments with different machine learning methods. (method) presented a superior performance, considering four metrics (accuracy, kappa index, sensibility and specificity).

This work is organized as following: in section 2 we present a brief of the state-of-the-art of DNA methods; in section 3 we present our methodology, including our proposal, the description of the database, the experiments parameters and the metrics used for performance measure. In section 4 we provide our experiments results and make analysis of them; finally, in section 6 we summarized the scientific contribution of this work and discusses the potential future work.

## 2. Related works

Several studies have sought to optimize the diagnosis of Covid-19 through the provision of rapid tests. The most common methods are based on the use of antibodies. Li et al. (2020) proposed a simple and rapid test for the combined detection of IgG and IgM antibodies. Both antibodies are indicative of infection. However, immunoglobulins M provide an immediate response to viral infections, and it can be detected in a period of 3 to 6 after infection. Immunoglobulin G, on the other hand, is important for the body’s long-term immunity or immune memory. With this in mind, they developed a test capable of detecting IgM and IgG simultaneously in blood samples, allowing detection in a longer time window. For the development of the rapid test, the authors collected samples from eight different laboratories and hospitals in China, with a total of 397 patient samples positive for Covid-19, and 128 negative samples. These results were confirmed by the RT-PCR technique using a respiratory tract specimen. Blood samples from patients were pipetted into the test kit, followed by two or three drops of dilution buffer. After 15 minutes it was possible to analyze the result using three markers. The first marker (letter C) or line on the display appears red when the sample is negative. The presence of IgG and IgM is indicated by red or pink lines in the regions with the letters M and G in the kit, and both antibodies may be present in the sample. The tone of the line is also indicative of the level of concentration of each type of antibody. Among the samples analyzed, the tests showed 88.66% sensitivity and 90.63% specificity. These values can be considered high, in comparison with results obtained in other studies (Cassaniti et al., 2020). The work also tested the performance of the method in 10 patients using peripheral blood. The results remained reliable. Thus, the work is promising and points out an interesting path for a simple and quick diagnosis, which can be an alternative for extensive testing of the population. However, the study does not point to tests with other types of viruses similar to SARS-Cov2, such as common flu. Given the similarity between viruses, the tests may indicate false positives, where the antibodies bind to similar antigens to SARS-Cov2.

Unlike rapid tests based on the detection of antigens, other works have sought to incorporate computational intelligence techniques in the diagnosis of Covid-19. Many of them have invested in automatic classification based on x-ray images making use of Deep Learning techniques, especially CNNs (Apostolopoulos & Mpesiana, 2020; Narin et al., 2020; Sethy & Behera, 2020). Apostolopoulos et al. (2020) applied these techniques to distinguish Covid-19 from other lung diseases, such as viral and bacterial pneumonias, pulmonary edema, pleural effusion, chronic obstructive disease, and pulmonary fibrosis. This study used a wide database with 3905 x-ray images, including approximately 450 cases of Covid-19. For model training, the images were scaled to 200×200 pixels. Small variations of the images were also considered. That is, the images were slightly rotated, in order to make the model robust to variations in position and orientation that may occur in the image acquisition process. To extract characteristics from the images, some models of convolutional networks (CNN) of the Mobile Net type were tested. Three techniques were compared: development of a new CNN architecture; application of a pre-trained CNN (Transfer Learning); and a hybrid method, which applies tuning strategies to specific layers of a pre-trained CNN. The experiments were carried out in Python, using the Keras library and TensorFlow as a backend. Among the tested configurations, the CNN developed from scratch showed the best results, suggesting that biomarkers related to Covid-19 can be found with the technique. The model achieved an average rating accuracy of 87.66%, considering all six classes. With special regard to Covid-19, the model achieved 99.18% accuracy, 97.36% sensitivity, and 99.42% specificity.

Gomes et al. (2020) also proposed the use of machine learning techniques for classification of x-ray images, distinguishing between Covid-19, viral pneumonia, bacterial pneumonia and healthy patients. In contrast to the previous work, the authors invested in low-cost computational methods. Thus, the authors tested Haralick and Zernike moments for extracting attributes and used classic classifiers, such as MLP, SVM, decision trees and Bayesian networks. The work points out that the chosen extractors can play an important role in the diagnosis by image. The reason for this is that in clinical practices it is common to find opaque and whitish areas in contexts of pneumonia. Finally, SVM reached the best performance. The authors reached an average accuracy of 89.78%, average recall and sensitivity of 0.8979, and average precision and specificity of 0.8985 and 0.9963 respectively. An initial desktop version of the system was developed and made available for free non-commercial use on Github.

On the other hand, other studies have invested in Covid-19 diagnostic methods through intelligent systems based on blood tests. Methods like this can be useful mainly in contexts of unavailable rapid tests, functioning as a patient screening process. For the development of the this work, Soares et al. (2020) used a database made available by the Israeli Hospital Albert Einstein, located in São Paulo, Brazil. The database has 108 clinical exams and data from 5644 patients. The authors chose 599 patients, who had few missing data (at least 16 tests performed). Among them, 81 had a positive result for Covid-19 by the RT-PCR method. In addition, they selected tests that can be performed quickly in an emergency context. The selected blood tests were complete blood count, creatinine, potassium, sodium, C-reactive protein, in addition to the patient’s age. Considering the imbalance of the database, the work used SMOTE techniques (Synthetic Minority Oversampling Technique) (Chawla et al., 2002; Lusa et al., 2013), which is capable of generating synthetic data from the minority class. Then they trained 10 support vector machines (SVM). The initial prediction model corresponds to the average probability of the 10 models developed. The testing and training processes were performed 100 times, using different subsets, with a 90% percentage split for training and 10% for testing. All models and statistics were obtained using R. The authors achieved an average specificity of 85.98%, an average sensitivity of 70.25%, a negative predictive value (NPV) of 94.92%, and a positive predictive value (PPV) of 44.96%. For the last metric, the authors believe that severe cases, however negative for Covid-19, generated more confusion in the classification. Another study Barbosa et al. (2020), using the same initial database, applied attribute extraction methods (Particle Swarm Optimization) to search for the best tests among the 108 initial ones. Then, the authors manually selected exams in order to reduce costs. The result was 24 selected exams, with performance similar to the initial base. The results of the evaluation metrics were: 95.16% of average accuracy, sensitivity of 0.969, specificity of 0.936 and 0.903 of kappa index. The authors made a desktop version of the system available for free non-commercial use.

While rapid diagnostic methods are important and optimize this process, the gold standard and recommendation of WHO is still the RT-PCR method with DNA sequencing (WHO, 2020a), similar to the method developed for the diagnosis of SARS-Cov (Chan et al., 2004; Emery et al., 2004; Corman et al., 2012). Thereby, multiple studies and protocols for identifying SARS-Cov2 by molecular diagnosis have already been published (Corman et al., 2020b,a; Poon et al., 2020; Chu et al., 2020; Nao et al., 2020). Chu et al. (2020) developed RT-PCR assays to detect SARS-Cov2 in human clinical samples. The authors relied on the first publication of the virus sequence on Genbank, in addition to sequences of other types of coronavirus to perform the alignment. Thus, they designed two monoplex assays, which target the ORF1b and N gene regions. Then, these primer and probe sequences were confirmed with other released SARS-Cov2 sequences. RT-PCR reactions were performed by a thermal cycler, using typical reaction mixture, forward and reverse primers, probe, and RNA sample. RNA and DNA purification kits were also used for extraction. The time for each RT-PCR run was about 1h and 15 min. In order to determine assays specificity, they used negative control samples with RNA extracted from other viruses (MERS, camel coronavirus, influenza A and B, adenovirus, enterovirus, rhinovirus, etc.) and from healthy patients. In contrast, all viruses belonging to the Sarbecovirus subgenus (SARS-like coronaviruses, and other coronaviruses) were considered positive in these assays. This decision was made due to the small amount of data available from SARS-Cov2 at the time of the development of the work. The study tested the method on two patients with suspected SARS-Cov2 infection. The samples were taken from different locations (sputum vs. throat swab) and at different infection periods (day 5 vs. day 3). Both patients received a positive result. Finally, the study results demonstrated the clinical value of respiratory samples for molecular diagnosis of Covid-19. The authors also observed a high sensitivity of the N gene for detecting the disease, being recommended as a screening assay, and the Orfb1 as a confirmatory one. The biggest difficulty, however, is that RT-PCR is time-consuming and labour intensive, and consequently, its result can take days to be available (Ai et al., 2020). This makes clinical conduct difficult and favors the contamination of more people by SARS-Cov2. In this sense, the objective of this work is to propose an optimization of the gold standard method.

## 3. Materials and methods

### 3.1. Proposed method

Our work considers genome sequences of several virus types, where each sequence is organized into a single matrix. Initially, the genome sequence is divided into *n* subsequences, which will then be overlapped with its neighbors. In the overlapping process, a parameter received by the method determines the size of the superimposed pieces. Every subsequence *i* is combined with a piece of the subsequence immediately to its left *i −* 1, and also with a piece of the one to its right, *i* + 1. An exception is made for the first and last sequence of the matrix, given that they have only one subsequence from which to take a piece. This procedure results in two new sequences for each of the subsequences generated from the original genome.

After that, these smaller sequences are represented by co-occurrence matrices. The matrices are square with 4*x*4 dimensions. Each element of the matrix represents the number of occurrences of a given pair of nucleotide bases, as well as the relationship between bases and their immediate neighbors. These elements are AA, AC, AT, AG, CA, CC, CT, CG, TA, TC, TT, TG, GA, GC, GT, and GG. The matrix is then normalized, where its maximum value is used to divide each of its elements.

Finally, all the 4 *×* 4 matrices are stacked together, forming a volume 4×4×*m*, wherein *m* is the number of subsequences resultant from the overlapping process. In general terms:

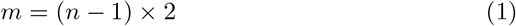

After obtaining this set of matrices, they are then concatenated, forming attribute vectors. These extracted attributes correspond to a high-level vectorial representation of the initial DNA sequence, independent from its size.

This process is illustrated in the following diagram in Figure 1.

**Figure 1:**
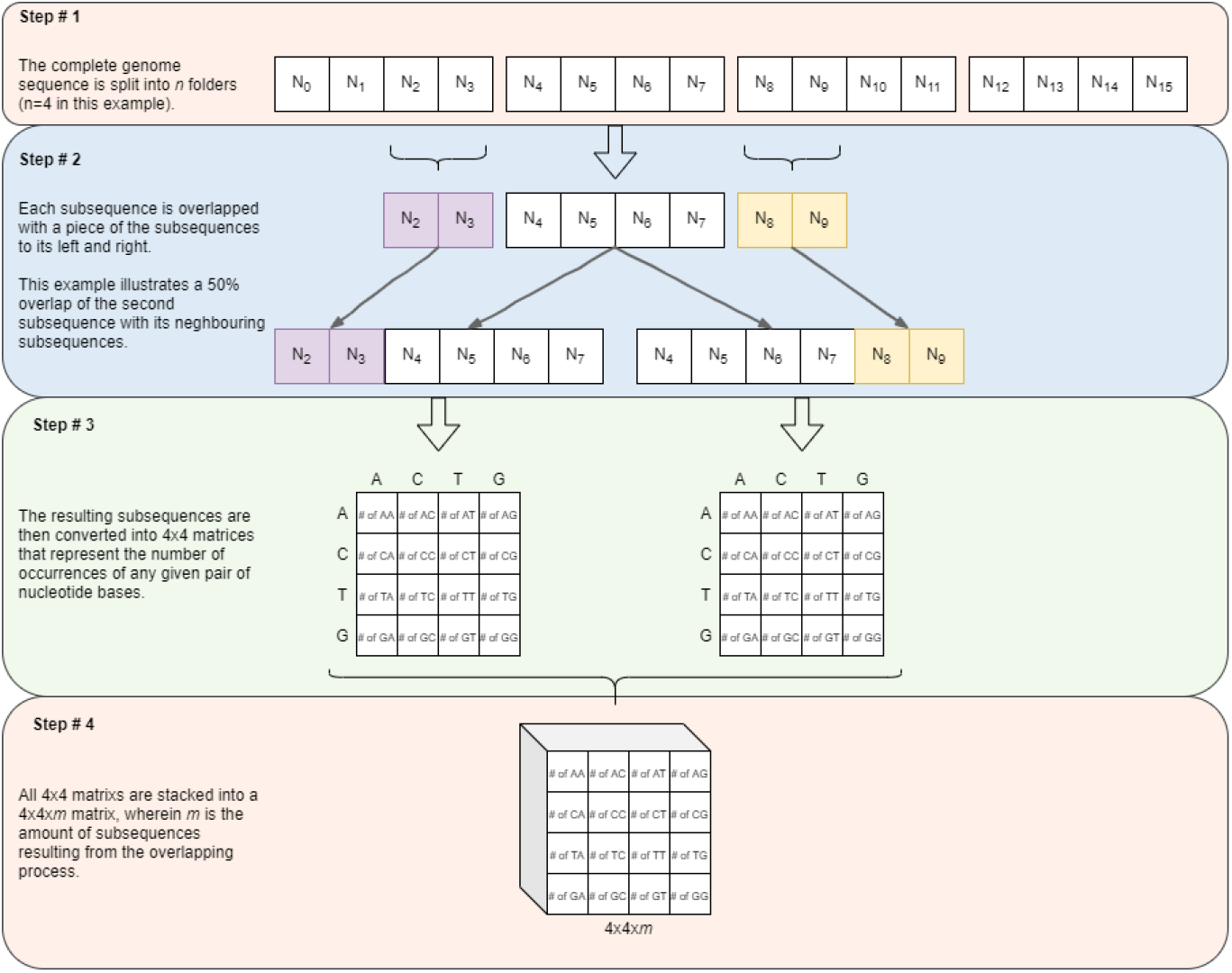
Steps of the proposed method: a new technique for representing genome sequences based on the analysis of the relationship between nitrogenous bases. It works as follows: the complete genome sequence is subdivided into *n* folders. Each subsequence is combined with a piece of its neighbors, generating two new sequences. These smaller sequences are represented by co-occurrence matrices, considering the occurrence of each of the nitrogenous bases, and the relationship between bases and their immediate neighbors. In the next step, theses matrices are stacked together as a volume. Finally, this set in concatenated, forming attribute vectors, which are a high-level vectorial representation of the original sequence.

### 3.2. Classifiers

In order to verify the proposed method’s efficiency in extracting characteristics from genome, different classifiers will process the data. The following classifiers were selected because they are widely used in machine learning.

#### 3.2.1. Random Forest

This classifier uses decision trees as its building blocks, Tin Kam Ho (1995). Decision trees, as illustrated in Figure 2, iteratively separate data by testing a property at a time, the resulting leafs representing the most specific category, and the root representing the raw data. The Random Forest is constructed of many such trees, that all have its own class prediction to any given input. The class with the most votes is the Random Forest’s output.

**Figure 2:**
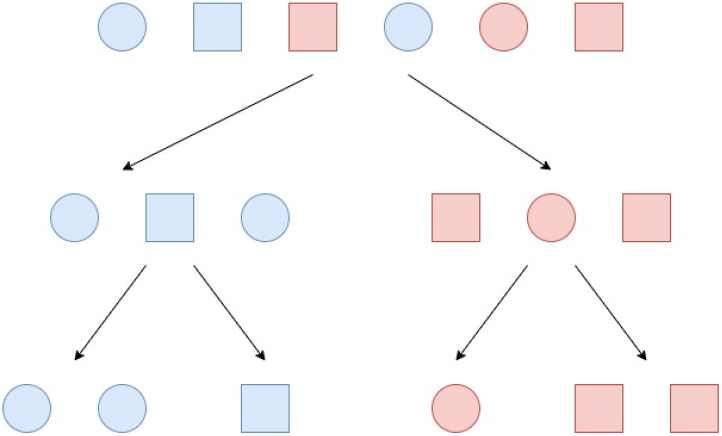
This decision tree example illustrates the classification of samples by two different features, colour and then shape.

As the characteristics that divide the genomes evaluated aren’t known, this method is advantageous because it verifies many possibly relevant properties. Thus, it can test and locate differences in the genetic code in question.

#### 3.2.2. Naive Bayes Classifier

This machine learning model uses probability, specifically the Bayes theorem, Maron (1961). The Bayes Theorem defines the probability of an event A happening, given that another event B has already taken place. The Bayes Theorem can be expressed as:

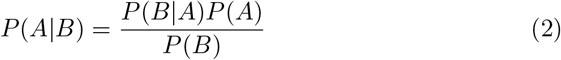

It is called naive because it assumes independence in the features that lead to the events. Furthermore, it assumes all predictors have an equal weight. This approach is beneficial because it explores the possibility that the genomes have dividing properties that are not correlated. Should that be the case, this classifier might yield good results.

#### 3.2.3. Instance Based Learner

This algorithm, also known as IBK, Altman (1992), doesn’t construct a model, but instead predicts by using a distance *k* between samples in the training set and a test sample. The training set instances selected generate the prediction, as demonstrated in Figure 3. It could prove to be successful, because it classifies by finding similar instances. Thus, it might be able to identify genome sequences that belong to the same virus.

**Figure 3:**
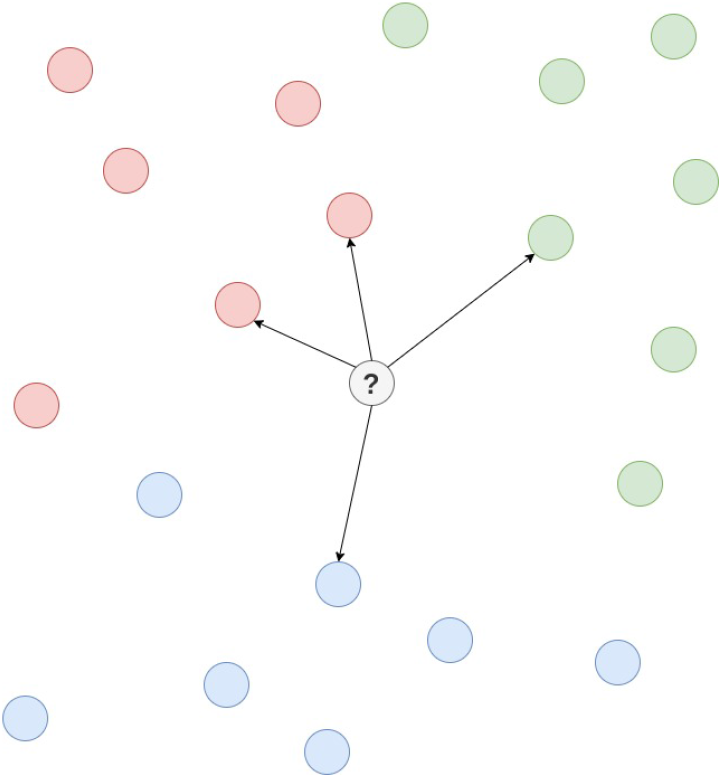

#### 3.2.4. Multilayer Perceptron

This classifier, shown in Figure 4, is a neural network capable of solving non linear data problems, Minsky & Papert (1969). Each neuron unit has weights that multiply the input, which is in turn processed by an activation function to generate the output. The weights are adjusted until the network can satisfy a certain accuracy in output. In this manner, it could identify the features that are particular to each class.

**Figure 4:**
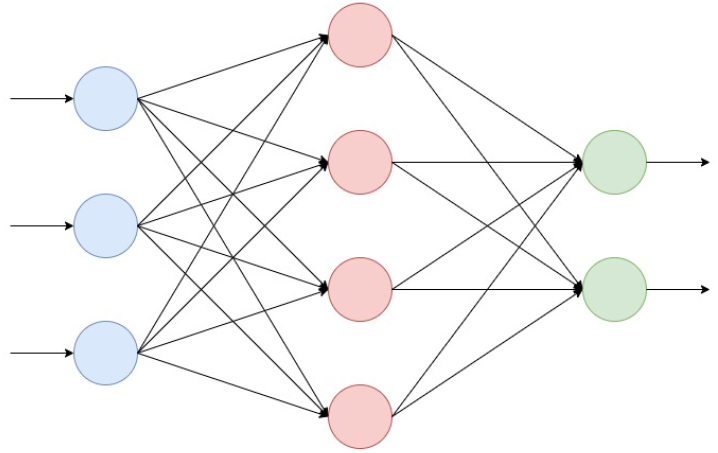
A multilayer perceptron with three layers of neurons.

#### 3.2.5. Support Vector Machine

This algorithm, Cortes & Vapnik (1995), hopes to find an optimal hyperplane that can separate the data into classes, as exemplified in Figure 5. The plane will have *n* dimensions, according to the number of features. The support vectors are the samples closest to the dividing hyperplane, that aid in its construction. Thus, it could be used to classify the genomes by dividing them with such a hyperplane.

**Figure 5:**
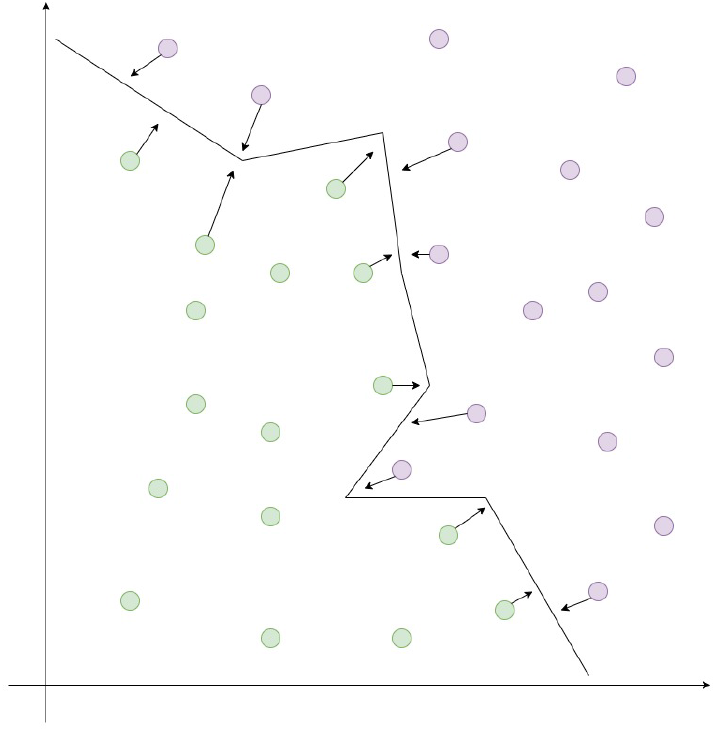
A binary classification problem, wherein the hyperplane created by the support vector machine has 2 dimensions.

### 3.3. Database

Twenty-five different viruses were used to evaluate the efficiency of the feature extraction method, including the SARS-Cov2 Cleemput et al. (2020). Data was obtained from the NIAID Virus Pathogen Database and Analysis Resource (ViPR) Pickett et al. (2012), which features multiple whole-genome sequences (WGS) from several viruses. Table 1 displays the number of examples per virus for each of the selected viruses.

**Table 1:** Number of instances in each class of virus.

| **Virus** | **Instances** |
| --- | --- |
| Phasmaviridae | 42 |
| **SARS-CoV-2** | **171** |
| Hepeviridae | 643 |
| Poxviridae | 697 |
| Ebola virus | 768 |
| Nairoviridae | 7977 |
| Filoviridae | 869 |
| Zika virus | 919 |
| Lassa virus | 1110 |
| Pneumoviridae | 1831 |
| Arenaviridae | 1840 |
| Togaviridae | 1983 |
| Caliciviridae | 2010 |
| Paramyxoviridae | 2609 |
| Rhabdoviridae | 2621 |
| Hantaviridae | 2785 |
| Phenuiviridae | 3089 |
| Peribunyaviridae | 3245 |
| Coronaviridae | 3256 |
| Enterovirus | 3784 |
| Dengue | 5885 |
| Picornaviridae | 5894 |
| Flaviridae | 14658 |
| Reoviridae | 62454 |
| Hepatitis C virus | 216223 |

The viruses have different sample sizes, ranging from 42, as is the case for Phasmaviridae, to 216,223, for Hepatitis C. The bar graphs below depict the distribution of sample sizes in both a linear and a logarithmic scale.

The second dataset used in this paper is from the Genome Reference Consortium Consortium (2013). Its purpose was to represent the human genome, and it has 103,959 samples.

### 3.4. Experiment setups

Various experiments were constructed to evaluate feature extraction method’s quality. They aim to simulate different use cases wherein SARS-CoV2 could need to be identified. There is a multiclass experiment, a binary classification, classification of viruses with similar symptoms and a real test scenario.

#### 3.4.1. Multiclass Classification

This experiment’s purpose is to differentiate SARS-CoV2 and the other viruses listed in table 1 from each other. In it, all 25 classes of the table 1 were used to build the database, that was split in training set and test set. In classes with more than 500 instances, the training set consisted of 500 them, and the rest were used in testing. The classes with less than a 500 samples had 70% of their samples allocated for training and 30% for testing. Additionally, the feature extraction hyperparameter *n* was set to 4, and overlap was tested at 30%, 50%, and 70%.

#### 3.4.2. Binary Classification

This test was utilized to analyze the proposed method’s efficiency in differentiating SARS-CoV2 from Coronaviridae. Viruses from the same family could potentially be challenging to classify when compared because they have a more similar genome. To account for that scenario, the two classes with their genomes are contrasted only to each other. Train and test splitting was performed exactly as in the multiclass evaluation. The feature extraction hyperparameter *n* was set to 4, and overlap was set to 30%, a percentage that was previously shown to represent the virus genome sequences satisfactorily.

#### 3.4.3. Viruses with similar symptoms

A third test was outlined to classify viruses with similar symptoms to SARS-CoV2. This should prove useful in determining if a patient has symptoms that indicate they might have SARS-CoV2, but other possibilities cannot be ruled out. Four classes were established: SARS-CoV2; Coronaviridae; Paramyxoviridae; Peneumoviridae, Hantaviridae, Enterovirus, and Nairoviridae. The train and test splits and the hyperparameter *n* were maintained as in previous tests. Overlap was set to 30%, 50%, and 70%.

#### 3.4.4. Real test scenario

This test included three classes: the human genome, from the Genome Reference Consortium Consortium (2013), SARS-CoV2 and the other viruses from table 1. It tests the real use case of the proposed method, wherein SARS-CoV2 must be identified amongst both human genome and other viruses. The train and test splitting was performed as previously established, and the value of *n* remained the same. Furthermore, the overlap was also tested at 30%, 50%, and 70%.

### 3.5. Metrics

- Confusion Matrix The confusion matrix provides a more straightforward structure for the portrayal of the model’s output, wherein the rows represent its predictions, and the columns represent the expected results. The confusion matrix layout used to display the results is illustrated in the following table, and its correct interpretation is as stated previously. Furthermore, *n* expresses the total number of instances, and each row, when summed, amounts to the total number of instances per class. The number of correctly classified instances can be obtained by adding all the elements in the main diagonal. On the other hand, the number of misclassified instances is obtained from the opposite diagonal.
- Accuracy The accuracy describes the rate of correct classification of instances and is the most commonly used metric in machine learning. Considering a confusion matrix *T* = [*t_i,j_*]_*n*×*n*_ for a classification task with *n* classes, in which *i* denotes the index of the *i*-th true class and *j* points to the index of the class associated to the classification decision, the *j*-th class, the accuracy is defined as following:

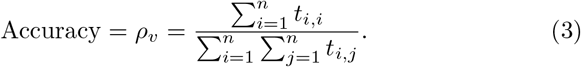
- Kappa Coefficient The Kappa Coefficient (*κ*) assesses the relation between the classified instances. It is defined as:

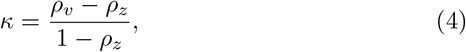

where

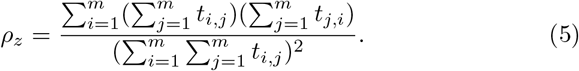
- Precision Precision indicates the proportion of positive and correct classification, and is thus calculated:

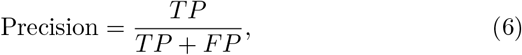

where TP is the number of true positives and FP is the amount of false positives.
- Recall Recall measures the proportion of actual positives correctly classified by the model. It is computed by:

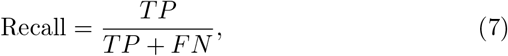

where FN is the number of false negatives.
- Sensitivity The sensitivity, or True Positive Rate, is given by:

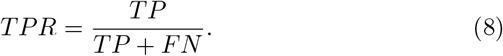
- Specificity The specificity, or True Negative Rate (TNR), if defined as following:

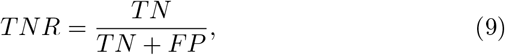

where TN is the number of true negatives.
- Area Under the ROC Curve The Receiver Operating Characteristic (ROC) curve is a graph that plots the True Positive Rate (TPR) and False Positive Rate (FPR) of classification for different thresholds. The FPR is defined by:

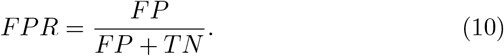 Thus, the Area Under the ROC Curve (AUC) measures performance for all possible thresholds of classification in a given model, and therefore it portrays the quality of results independently of it.

## 4. Results

### 4.1. Multiclass Classification

In order to evaluate the efficiency of the proposed features extraction method, this first round of experiments was conducted in a more challenging scenario with twenty-five different viruses, including the SARS-CoV2. Five types of classifiers were tested: IBk, Multilayer Perceptron (MLP), Naive Bayes classifier (NBC), Random Forest, and Support Vector Machines (SVM). All experiments were performed with Weka software. The parameters used in each machine learning method is shown in Table 2.

**Table 2:** Classifiers parameters: SVMs with linear kernel; MLPs with 48 neurons in the hidden layer; random forests with 100 trees; and standard IBK and Bayesian networks.

| Classifier | Hyperparameters |
| --- | --- |
| Random Forest (RF) | Number of estimators: 100 |
| Naive Bayes Classifier (NBC) | – |
| IBK | Number of neighbors to use: 1<br>Distance metric: Euclidean distance |
| Multilayer Perceptron (MLP) | Learning rate: 0.3<br>Momentum: 0.2<br>Single hidden layer with 48 neurons (number attributes divided by two)<br>Sigmoid activation function |
| Support Vector Machine (SVM) | C: 0.1<br>Linear Kernel |

Figure 6 shows the accuracy for all classifiers in the datasets with 30%, 50%, and 70% of overlap, respectively. Considering this multiclass classification, all three datasets (with 30%, 50%, and 70% overlap) presented Random Forest classifier with the highest accuracies (approximately 94% in all the datasets).

**Figure 6:**
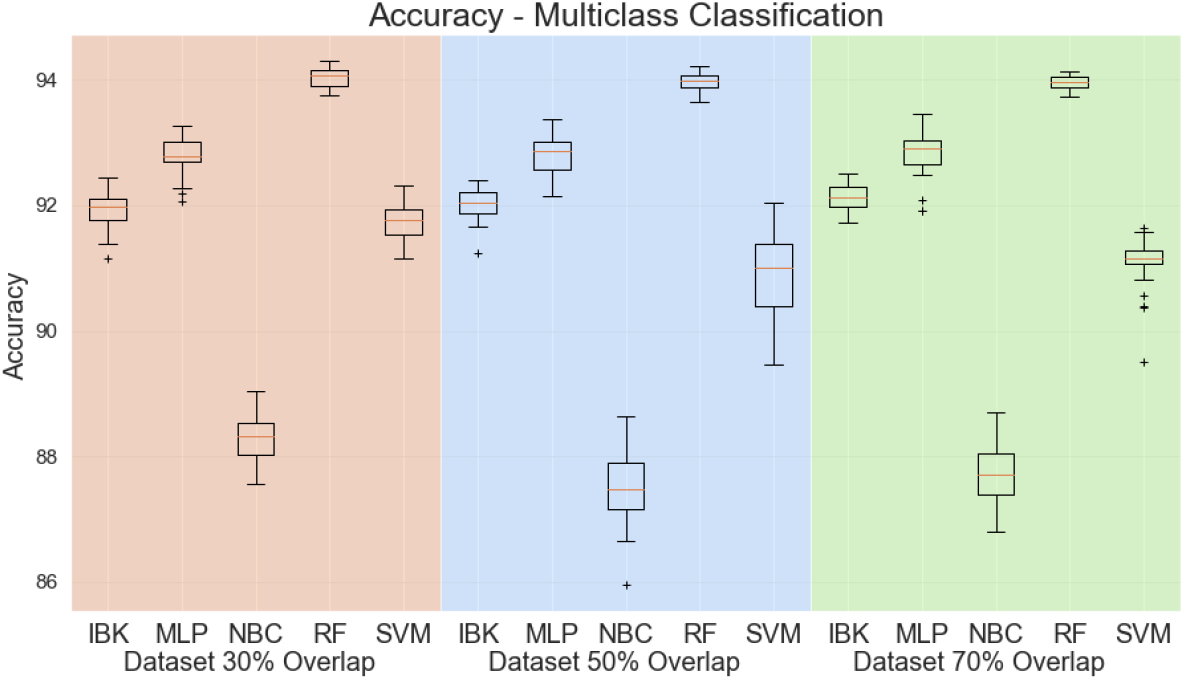
Accuracy for multiclass scenario.

Figure 7 shows box plots for the Kappa statistic. Since Kappa statistic is less sensitive to the high imbalanced test dataset, it is a better evaluation metric then accuracy. Nevertheless, the Random Forest classifier achieves the highest Kappa statistics compared with the other classifiers (above 0.88 in all experiments).

**Figure 7:**
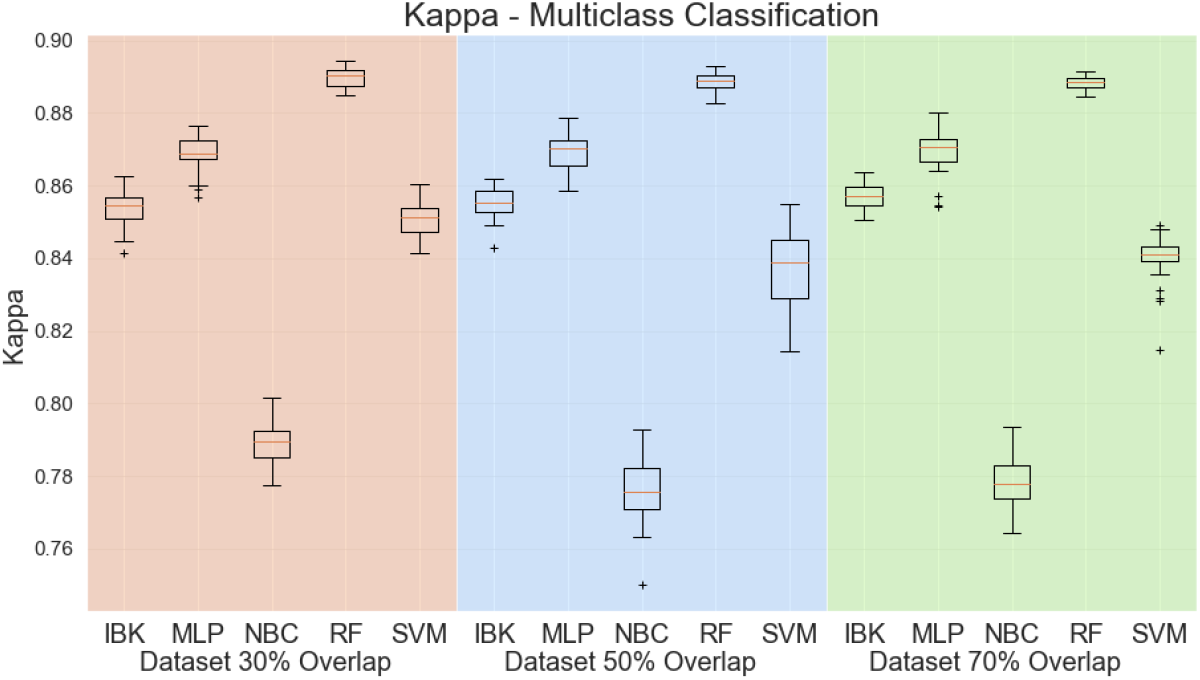
Kappa statistic for multiclass scenario.

Besides, accuracy and Kappa statistic, Figure 8 shows the weighted average sensitivity, specificity, and ROC area for all datasets and classifiers. For the weighted average sensitivity and ROC area, Random Forest results are higher or equal to other classifiers. For the weighted average specificity, visual analysis of Figure 8-b suggests that the IBK classifier achieves higher scores on this metric. However, all classifiers, except Naive Bayes Classifier, achieved results above 0.99 on weighted average specificity, so the Random Forest is presented as a robust classifier for this task.

**Figure 8:**
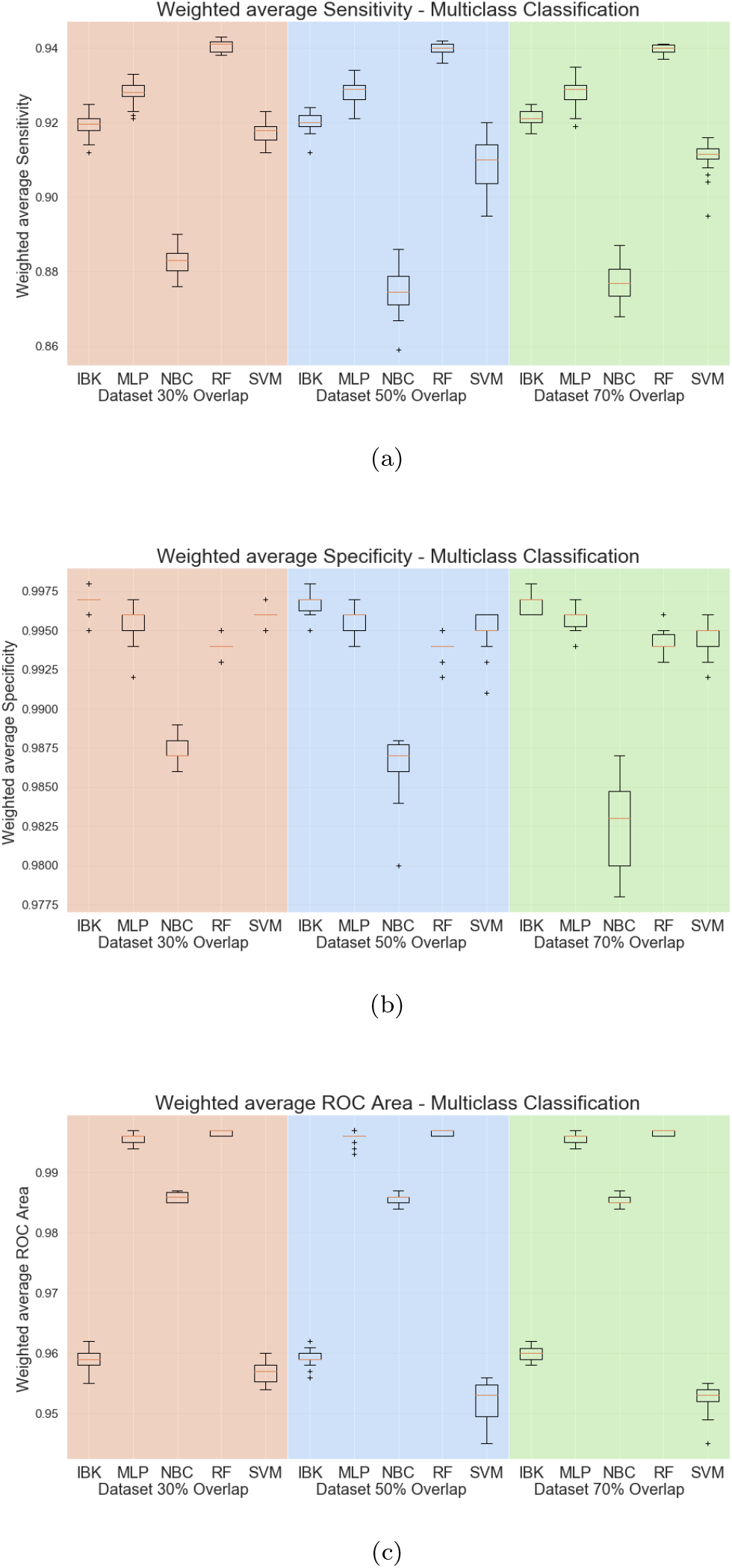
Weighted average sensitivity (a), specificity (b), and ROC area for Multiclass test scenario.

Aiming to evaluate the overlap percentage in the feature extraction method, Figure 9 shows box plots for accuracy, Kappa statistic, weighted average precision, recall and ROC area for the Random Forest classifier in the datasets with 30%, 50%, and 70% overlap percentages. The variance of accuracy and kappa in the dataset with 30% overlap is higher than in the 50% and 70% overlap dataset. However, 30% overlap seems to be slightly better (or at least at the same level) as the others overlap percentages.

**Figure 9:**
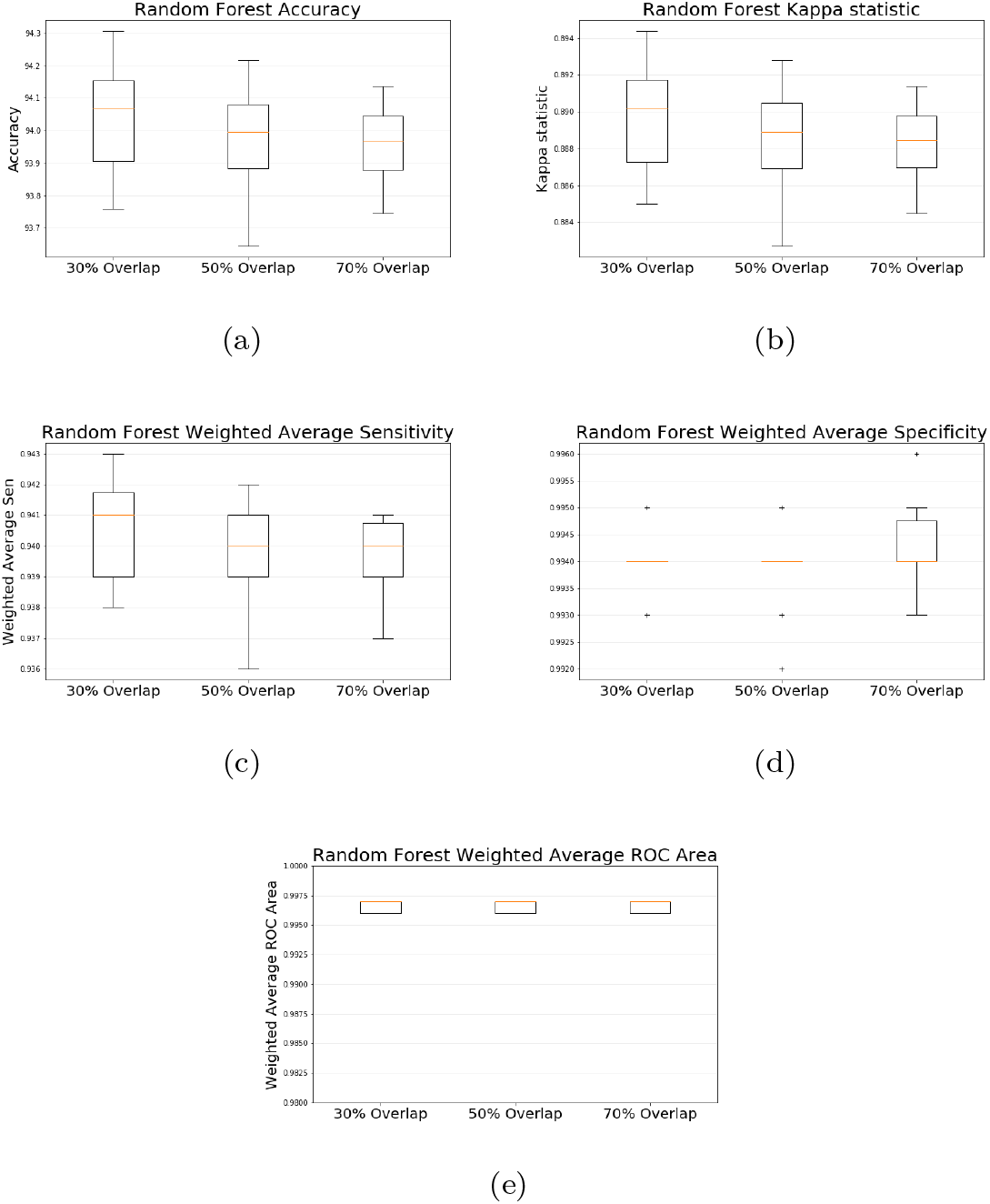
Random Forest accuracy (a), kappa (b), weighed average sensitivity (c), specificity (d) and ROC area (e) in 30%, 50% and 70% overlap percentages.

Because of class unbalancing in the test dataset, we need to evaluate sensitivity, specificity, and ROC area for each class individually. Considering the Random Forest classifier in the dataset with 30% overlap, Table 3 shows the results of sensitivity, specificity, and ROC area individually for each virus in the database. Specificity and ROC Area results are above 0.9 for every virus. The sensitivity varies from 0.99391 for Pneumoviridae to 0.23397 for Filoriviridae. However, for most of the classes, sensitivity has values greater than 0.8 (including SARS-Cov2 class with a sensitivity of 0.82).

**Table 3:** Random Forest sensitivity, specificity, and ROC area for every single class (results from dataset with 30% overlap).

| Class | Sensitivity-Recall |  | Specificity |  | ROC Area |  |
| --- | --- | --- | --- | --- | --- | --- |
|  | Average | Std. Dev. | Average | Std. Dev. | Average | Std. Dev. |
| Class 0 - Picornaviridae | 0.46387 | 0.01407 | 0.99674 | 0.00028 | 0.99223 | 0.00163 |
| Class 1 - Arenaviridae | 0.44609 | 0.01301 | 0.99906 | 0.00008 | 0.99773 | 0.00121 |
| Class 2 - Caliciviridae | 0.98755 | 0.00392 | 0.99886 | 0.00045 | 0.99893 | 0.00089 |
| Class 3 - Pneumoviridae | 0.99391 | 0.00285 | 0.99999 | 0.00001 | 0.99993 | 0.00025 |
| Class 4 - Phenuiviridae | 0.94210 | 0.00704 | 0.99966 | 0.00005 | 0.99840 | 0.00076 |
| Class 5 - Togaviridae | 0.99159 | 0.00327 | 0.99958 | 0.00016 | 0.99950 | 0.00067 |
| Class 6 - Poxviridae | 0.99314 | 0.00662 | 0.99994 | 0.00003 | 0.99947 | 0.00191 |
| Class 7 - Filoviridae | 0.23397 | 0.03284 | 0.99950 | 0.00005 | 0.99877 | 0.00096 |
| Class 8 - Flaviridae | 0.55250 | 0.00768 | 0.98242 | 0.00144 | 0.95860 | 0.00308 |
| Class 9 - Hantaviridae | 0.98284 | 0.00565 | 0.99960 | 0.00004 | 0.99907 | 0.00100 |
| Class 10 - Lassa virus | 0.59120 | 0.03118 | 0.99778 | 0.00006 | 0.99707 | 0.00112 |
| Class 11 - Dengue | 0.87925 | 0.01729 | 0.98384 | 0.00029 | 0.98910 | 0.00070 |
| Class 12 - Hepeviridae | 0.98924 | 0.00711 | 0.99972 | 0.00014 | 0.99933 | 0.00083 |
| Class 13 - Ebola virus | 0.39652 | 0.05417 | 0.99913 | 0.00004 | 0.99833 | 0.00094 |
| Class 14 - Enterovirus | 0.70652 | 0.02068 | 0.99144 | 0.00021 | 0.99313 | 0.00034 |
| Class 15 - Zika virus | 0.96675 | 0.01076 | 0.99728 | 0.00002 | 0.99800 | 0.00000 |
| Class 16 - Nairoviridae | 0.98687 | 0.00686 | 0.99974 | 0.00008 | 0.99843 | 0.00238 |
| Class 17 - Coronaviridae | 0.96194 | 0.00181 | 0.99988 | 0.00005 | 0.99833 | 0.00119 |
| Class 18 - Paramyxoviridae | 0.98781 | 0.00274 | 0.99982 | 0.00006 | 0.99970 | 0.00078 |
| Class 19 - Rhabdoviridae | 0.97468 | 0.00541 | 0.99976 | 0.00009 | 0.99967 | 0.00047 |
| Class 20 - Hepatitis C virus | 0.97425 | 0.00244 | 0.99288 | 0.00074 | 0.99823 | 0.00062 |
| Class 21 - Peribunyaviridae | 0.95344 | 0.00603 | 0.99870 | 0.00046 | 0.99523 | 0.00138 |
| Class 22 - Reoviridae | 0.98059 | 0.00261 | 0.99923 | 0.00014 | 0.99963 | 0.00048 |
| Class 23 - Phasma Viridae | 0.29167 | 0.10704 | 0.99999 | 0.00000 | 0.99000 | 0.01464 |
| **Class 24 - SARS-Cov2** | **0.82222** | **0.05613** | **0.99974** | **0.00001** | **0.99883** | **0.00250** |

In order to perform a visual analysis of these results, Figure 10 shows the average confusion matrix for the Random Forest classifier in the dataset with 30% overlap. The confusion matrix is expressed in terms of percentage for the particular class, and the classes indexes numbers are the same as shown in Table 3. We can see that for some classes, there is a confusion with another virus. For example, most of the Picornaviridae virus (index 0) is classified as Enterovirus (index 14). This confusion is not symmetrical: Picornaviridae is misclassified as Enterovirus, but Enterovirus is not misclassified as Picornaviridae. The only exception for this observation of confusion with another virus type is the Phasma Viridae (index 23), which is confused with two other viruses: Hantaviridae (index 9), and Peribunyaviridae (index 21). However, since there are few examples of Phasma Viridae in the dataset (only 42 examples), those results may be caused by the low representative of this class in the dataset.

**Figure 10:**
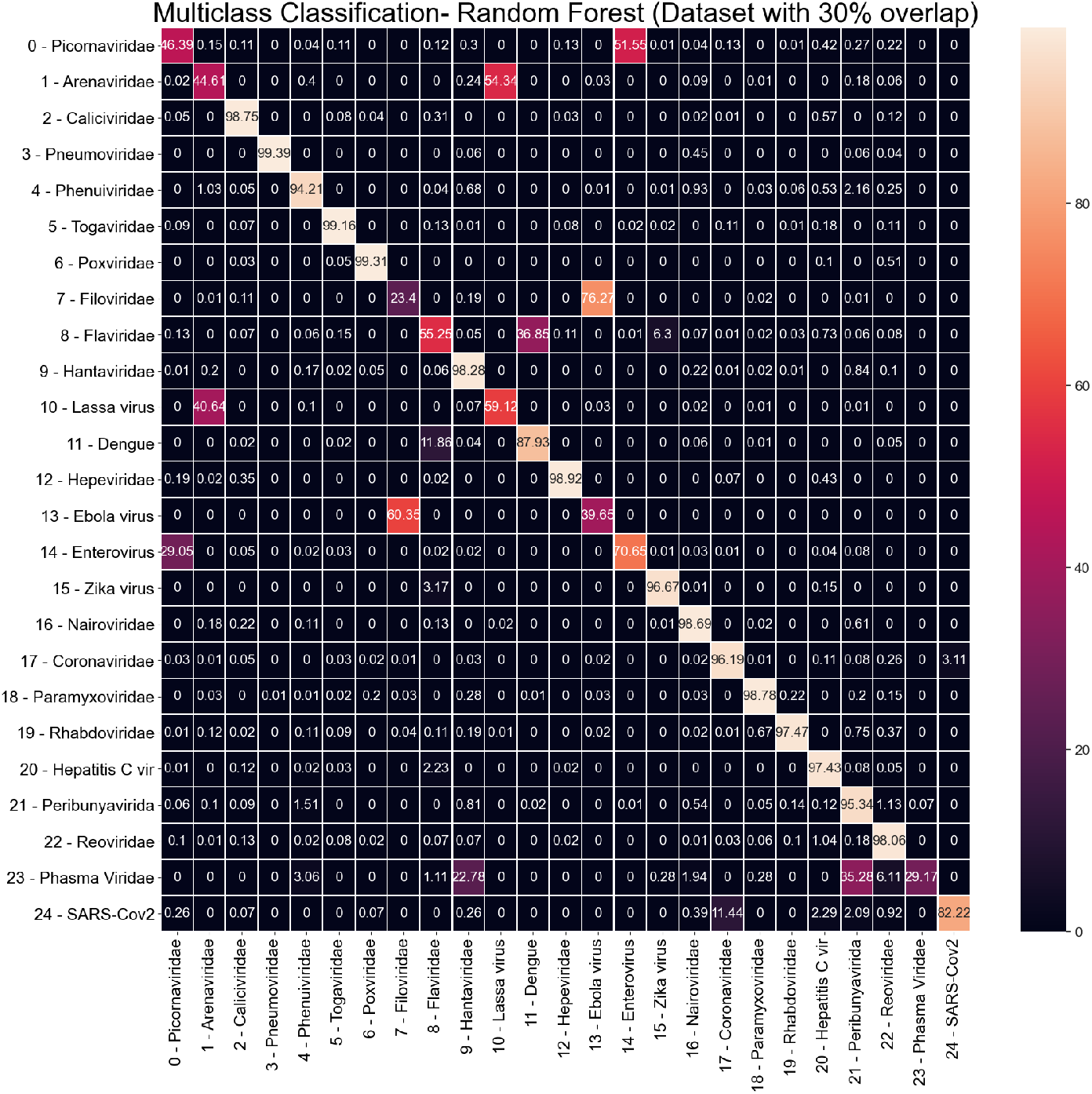
Random Forest average Confusion Matrix (results from dataset with 30% overlap).

Regarding the SARS-Cov2 virus (index 24), the only relevant confusion is with Coronaviridae (index 17). It is a predictable outcome since SARS-Cov2 belongs to the Coronaviridae virus family. 3.1% of Coronaviridae examples are classified as SARS-Cov2 (the only confusion noticed in column 24 of the confusion matrix). A more significant confusion is noticed between SARS-Cov2 and Coronaviridae since 11% of SARS-Cov2 are misclassified as Coronaviridae.

Since the ROC area for SARS-Cov2 is 0.99883 (Table 3), we performed a threshold adjustment for SARS-Cov2 class in order to reach 100% sensitivity. The new average confusion matrix is shown in Figure 11. Higher false positives for SARS-Cov2 remains from Coronaviridae (5,1% - index 17). In the sequence of false positive rates, we have: Hepatitis C virus (3,47% - index 20), Reoviridae (3,19% - index 22), and Phasma Viridae (2,68% index 23).

**Figure 11:**
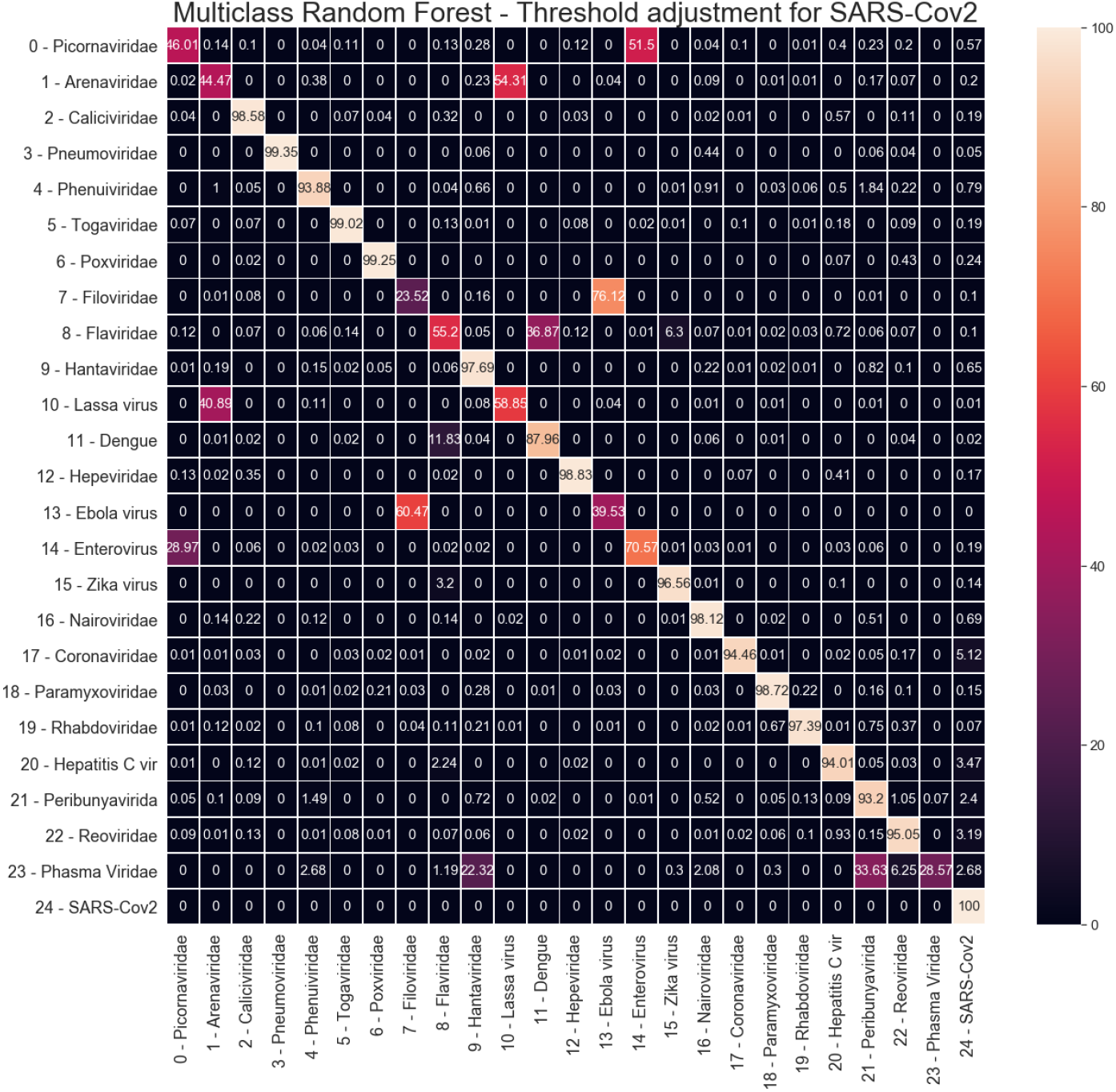
Random Forest average Confusion Matrix with threshold adjustment for 100% sensitivity on SARS-Cov2 (index 24).

### 4.2. Binary Classification

Given that, in the multiclass scenario, the highest false positives for SARS-Cov2 are from Coronaviridae, we evaluated the same classifiers used in the multiclass scenario for a binary classification between Coronavirus and SARS-Cov2. For this experiment, only the dataset with 30% overlap was used, since this overlap percentage has shown to represent the virus genome sequences satisfactorily.

Figure 12 shows the accuracy, kappa statistic, weighted average sensitivity, specificity, and ROC area for each classifier. It is important to state that there is still a relevant imbalance between the number of Coronaviridae and SARS-Cov2 examples in the dataset (3256 and 171, respectively). So, the Kappa statistic is still more appropriate than accuracy to assess the classifier’s overall performance. Regarding Kappa statistics, weighted average specificity, and ROC area, MLP results are higher or equal to other classifiers. For the weighted average sensitivity, SVM achieves higher results than MLP. Nevertheless, given that average sensitivity for MLP is higher than 0.96 and MLP overcomes SVM in all other metrics, MLP seems to be a more robust classifier for this particular task.

**Figure 12:**
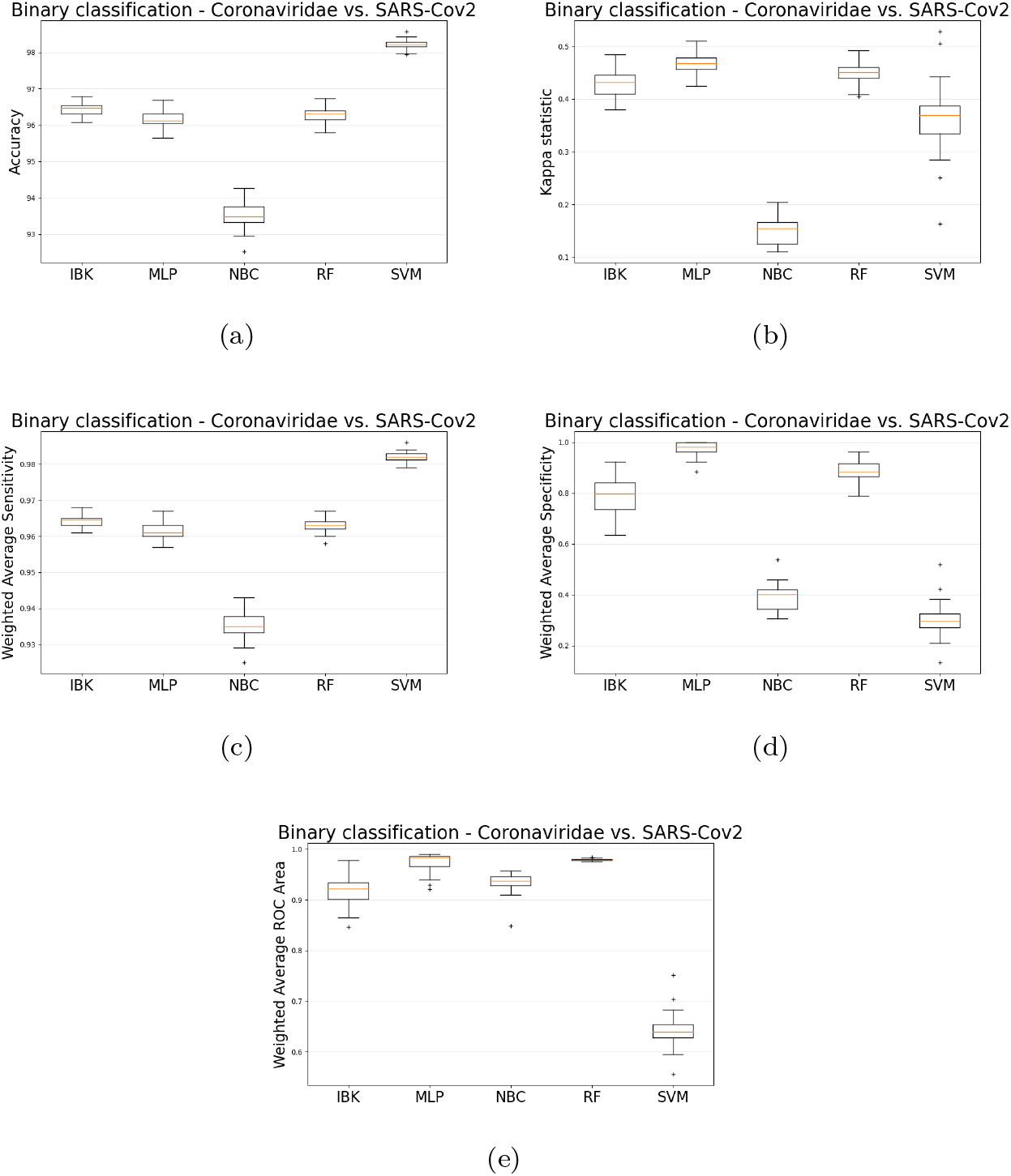
Binary classification (Coronavuris vs. SARS-Cov2 using the 30% overlap dataset) accuracy (a), kappa (b), weighed average sensitivity (c), specificity (d), and ROC area (e).

Table 4 shows the sensitivity, specificity and ROC Area for each class. It is possible to notice that each one of those metrics has values above 0.96. Figure 13 shows the average Confusion Matrix for MLP classifier. There was no relevant difference with the multiclass scenario regarding the confusion between Coronaviridae and SARS-Cov2 since there is still a 3.85% of Coronaviridae examples misclassified as SARS-Cov2. However, about the confusion between SARS-Cov2 and Coronaviridae, the binary MLP classifier achieved 2.61% of confusion while 11% in the multiclass scenario.

**Table 4:** Results of Sensitivity, specificity, and ROC area for MLP binary classifier (Coronavuris vs. SARS-Cov2 using the 30% overlap dataset).

| Class | Sensitivity-Recall |  | Specificity |  | ROC Area |  |
| --- | --- | --- | --- | --- | --- | --- |
|  | Average | Std. Dev. | Average | Std. Dev. | Average | Std. Dev. |
| Class 17 - Coronaviridae | 0.96151 | 0.00246 | 0.97386 | 0.03052 | 0.97353 | 0.01863 |
| **Class 24 - SARS-Cov2** | **0.97386** | **0.03052** | **0.96151** | **0.00246** | **0.97353** | **0.01863** |

**Figure 13:**
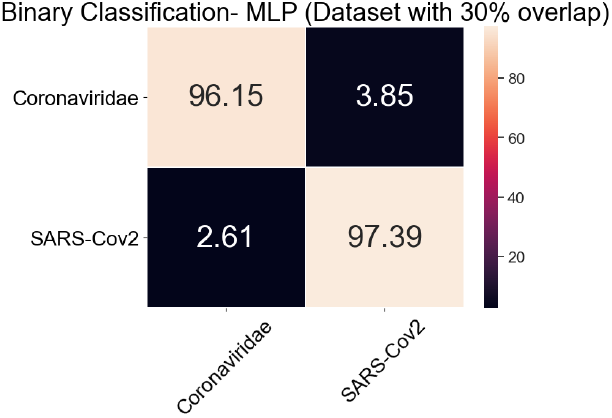
MLP average Confusion Matrix for binary classification task (Coronavuris vs. SARS-Cov2 using the 30% overlap dataset).

### 4.3. Viruses with similar symptoms

In this experiment, viruses were selected due to similar symptoms. The dataset was arranged into four classes: SARS-Cov2, Coronaviridae, Paramyxoviridae, and Miscellaneous. The Miscellaneous Class is a compound of Peneumoviridae, Hantaviridae, Enterovirus, and Nairoviridae. Then, the same classifiers used previously were evaluated in this classification task.

Figures 14 and 15 shows the accuracy and kappa for all classifiers and datasets in this classification task. Except for the Naive Bayesian classifier, classifiers have similar performance metrics, with approximately 97% accuracy and kappa equal to 0.96. Figure 16 shows the weighted average specificity and sensitivity and ROC are. The weighted average sensitivity and specificity look very similar to all classifiers (except Naive Bayes Classifier). However, the weighted average ROC area for MLP and Random Forest classifiers is slightly higher than the other classifiers, although IBK and SVM classifiers also achieve a weighted average ROC area above 0.98 in all datasets.

**Figure 14:**
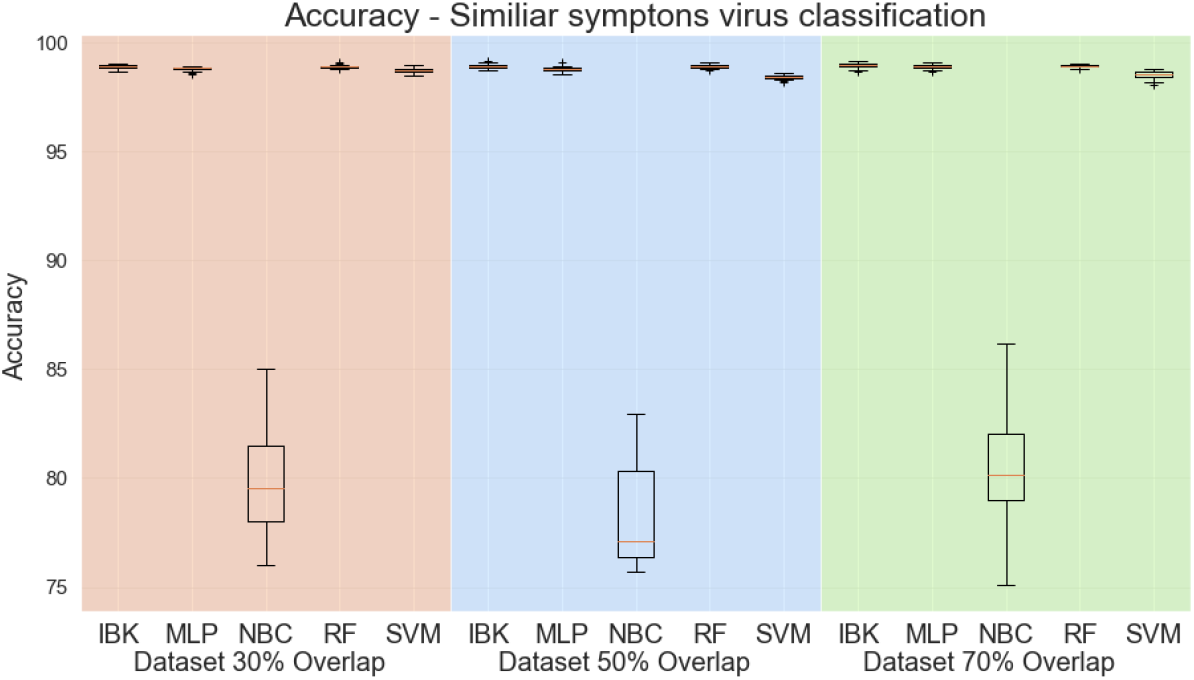
Accuracy for similar symptoms scenario.

**Figure 15:**
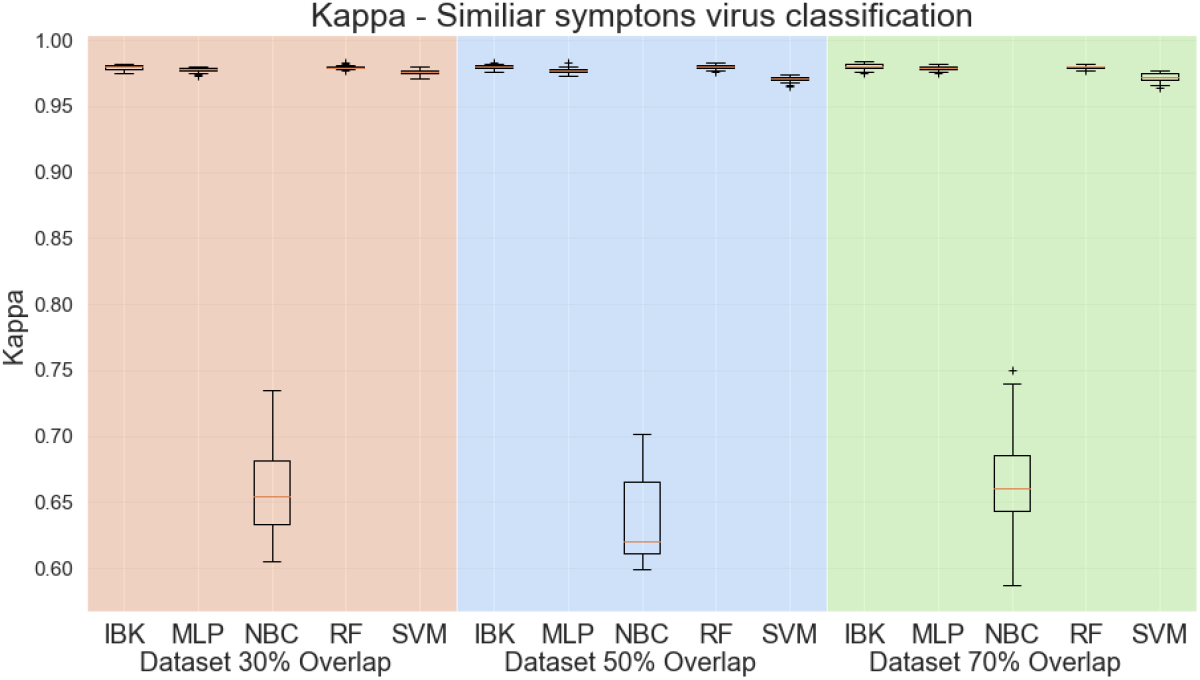
Kappa Statistic for similar symptoms scenario.

**Figure 16:**
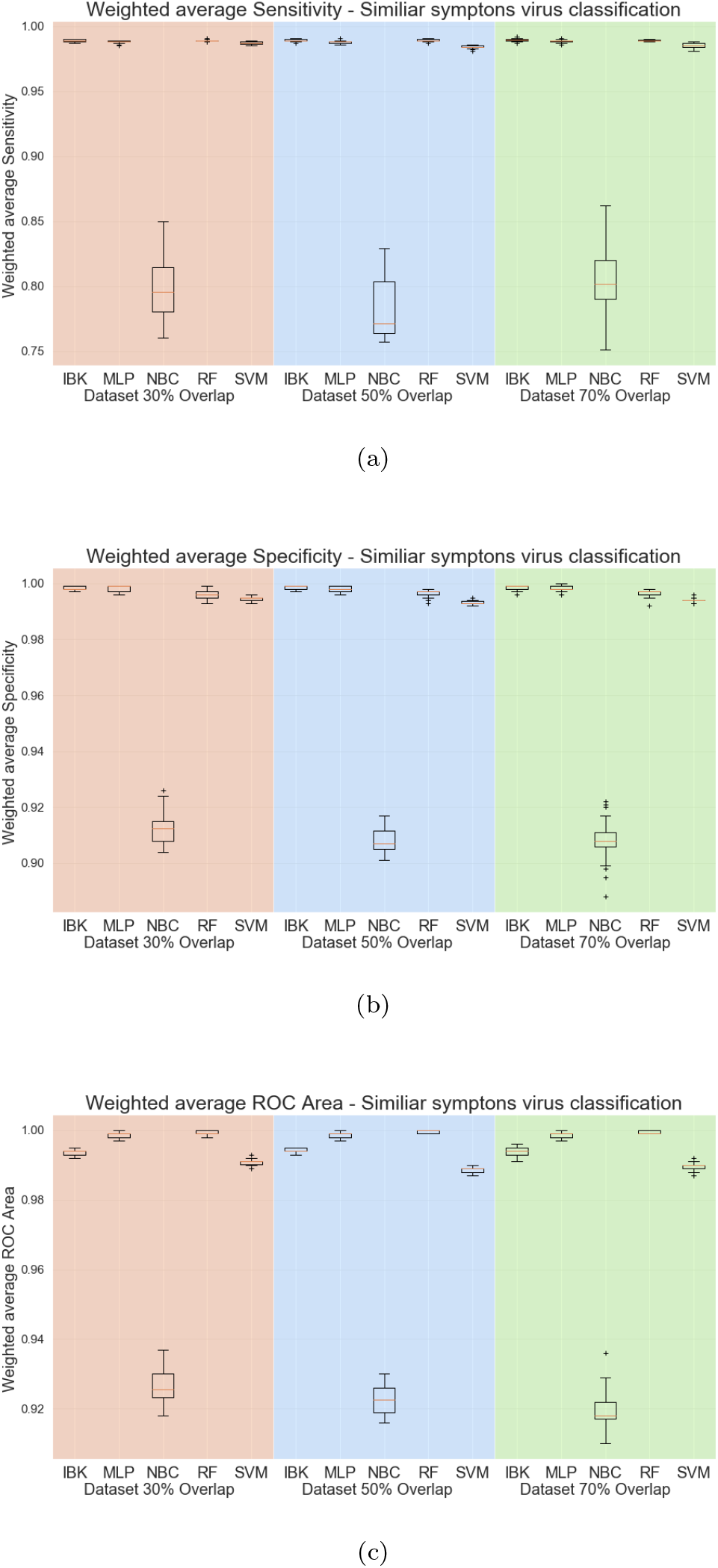
Weighted average sensitivity (a), specificity (b), and ROC area for Similar symptoms viruses test scenario.

In order to better evaluate the MLP and Random Classifier, Figure 17 shows the confusion matrices for those classifiers in all datasets. The Random Forest presents a confusion between the SARS-Cov2 and the Coronaviridae of approximately 10%. It is very similar to the achieved results in the multiclass scenario. However, the MLP classifier achieves significantly low-level confusions between SARS-Cov2 and Coronaviridae (1.57% in the datasets with 30% and 50% overlap). The main confusion found in the MLP classifier is between Conronarividae and SARS-Cov2 (3.81% for the dataset with 30% overlap). By MLP confusion matrix analysis is not possible to find significant differences between the 30%, 50%, or 70% overlap percentages. Since the 30% overlap requires less computational effort to extract the features, we can select the MLP classifier with a 30% overlap dataset as a better approach to this particular task. The Table 5 shows the sensitivity, specificity and ROC area for each class. The average ROC Area and specificity are above 0.99 for all classes. The average sensitivity is also above 0.99 for the Paramyxoviridae and Miscellaneous classes. The lowers sensitivity is for Coronaviridae (0.959), while a slightly higher sensitivity is achieved for SARS-Cov2 (0.97).

**Figure 17:**
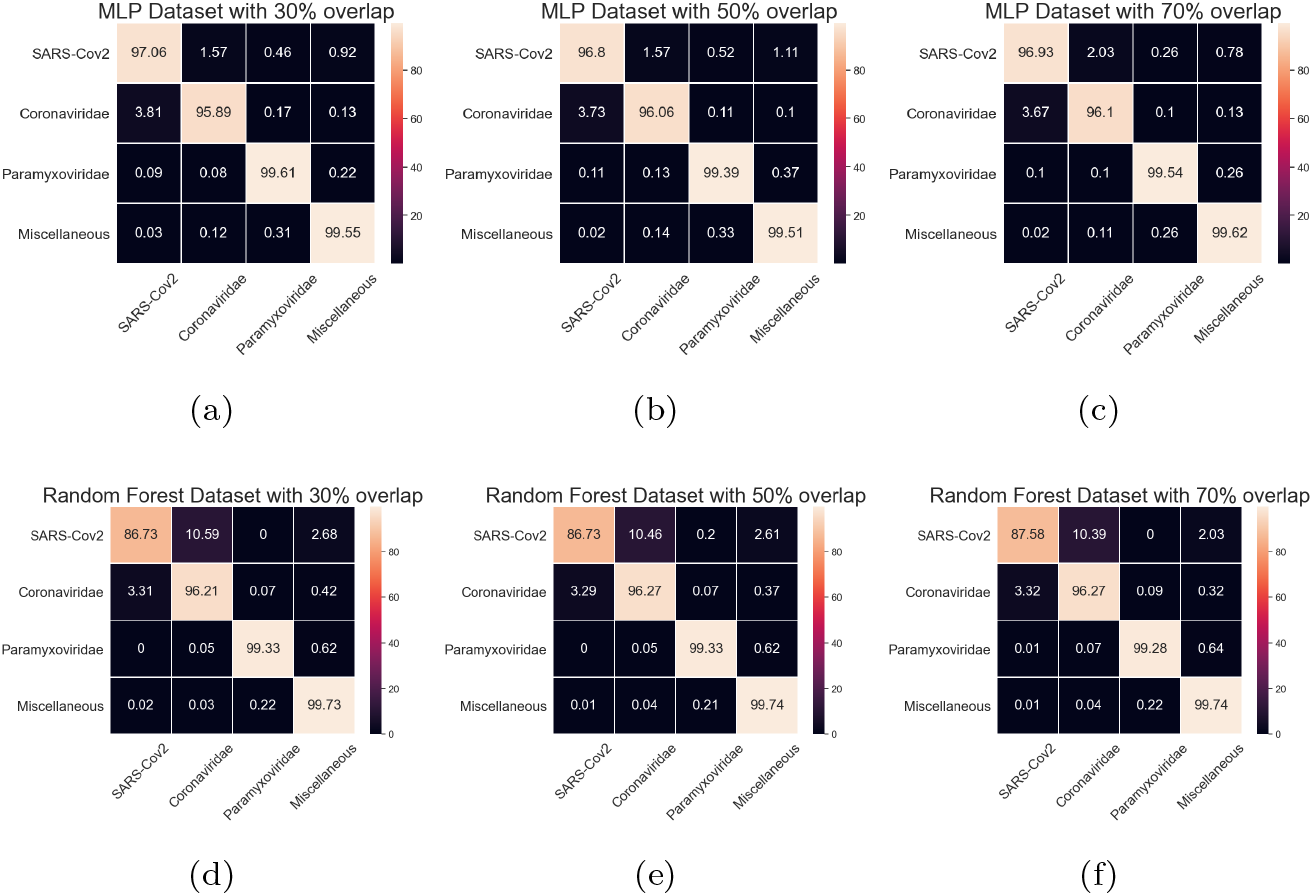
Average Confusion matrices for MLP and Random Forest in the Similar symptoms viruses test scenario.

**Table 5:** Results of Sensitivity, specificity, and ROC area for MLP classifier in the similar symptoms viruses test scenario (results from dataset with 30% overlap).

| Class | Sensitivity-Recall |  | Specificity |  | ROC Area |  |
| --- | --- | --- | --- | --- | --- | --- |
|  | Average | Std. Dev. | Average | Std. Dev. | Average | Std. Dev. |
| **SARS-Cov2** | **0.97059** | **0.03387** | **0.99187** | **0.00046** | **0.99583** | **0.00481** |
| Coronaviridae | 0.95891 | 0.00249 | 0.99882 | 0.00069 | 0.99687 | 0.00076 |
| Paramyxoviridae | 0.99611 | 0.00430 | 0.99726 | 0.00104 | 0.99863 | 0.00244 |
| Miscellaneous* | 0.99548 | 0.00151 | 0.99827 | 0.00153 | 0.99943 | 0.00072 |

\*Miscellaneous class includes four virus types: Pneumoviridae, Hantaviridae, Enterovirus, and Nairoviridae

### 4.4. Real test scenario

In this scenario, the SARS-Cov2 test is designed as a three-class classification problem: SARS-Cov2 (the test target), GRCh38 (the healthy human reference), and Coronaviridae (a virus control sample). The same classifies used in the other experiments were applied to this new task.

Figure 18 shows the accuracy and Figure 19 shows the kappa statistic results. Except for the Naive Bayes Classifier, all other classifiers have accuracy above 99% kappa above 0.9. By these metrics, It is not possible to distinguish the best classifier. The same behavior is observed in the weighted average metrics shown in Figure 20. Weighted average sensitivity, specificity, and ROC area are higher than 0.99 for all classifiers except the Naive Bayes Classifier.

**Figure 18:**
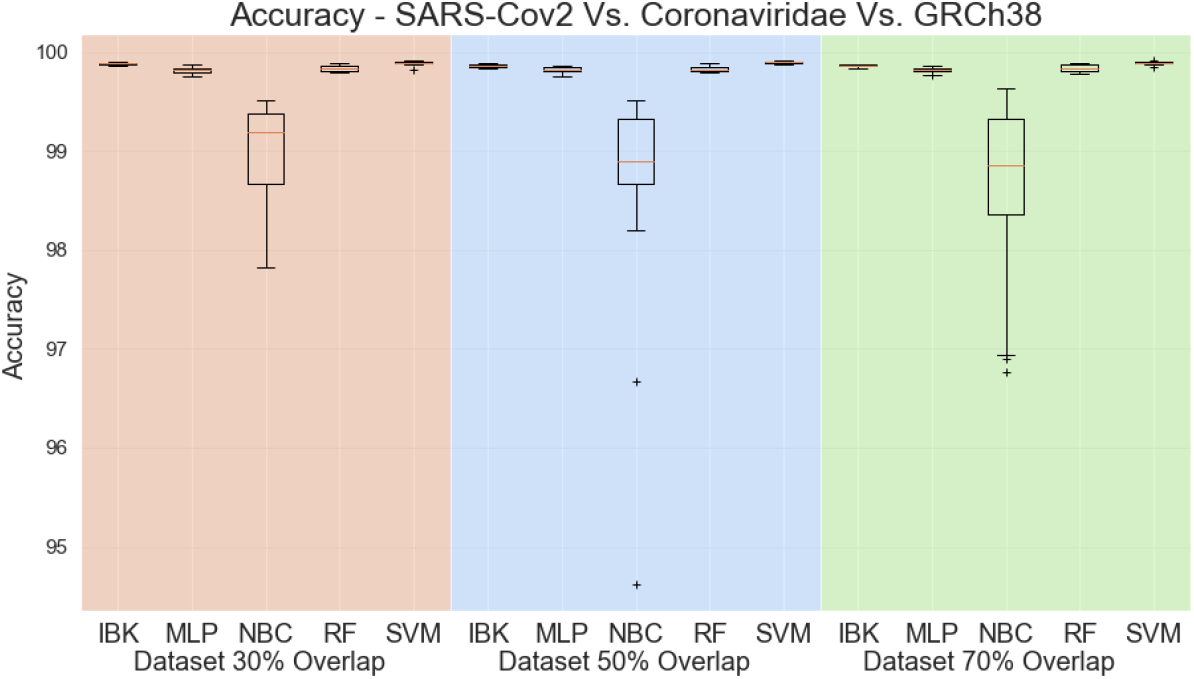
Accuracy for SARS-Cov2 test scenario.

**Figure 19:**
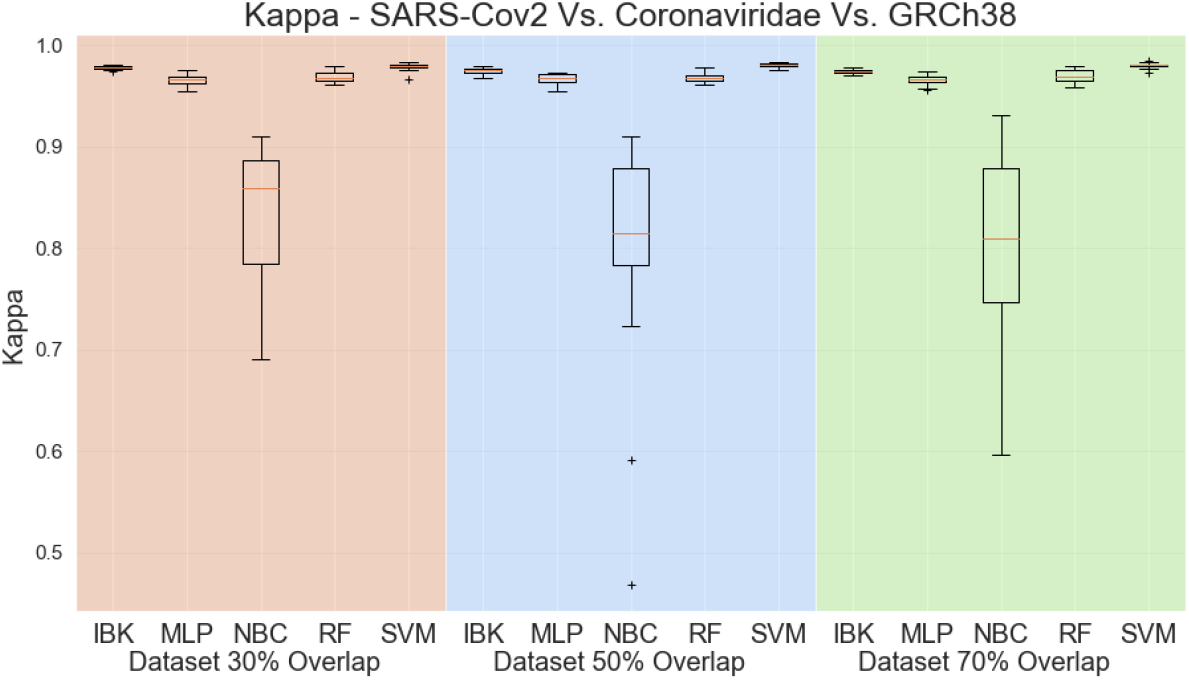
Kappa Statistic for SARS-Cov2 test scenario.

**Figure 20:**
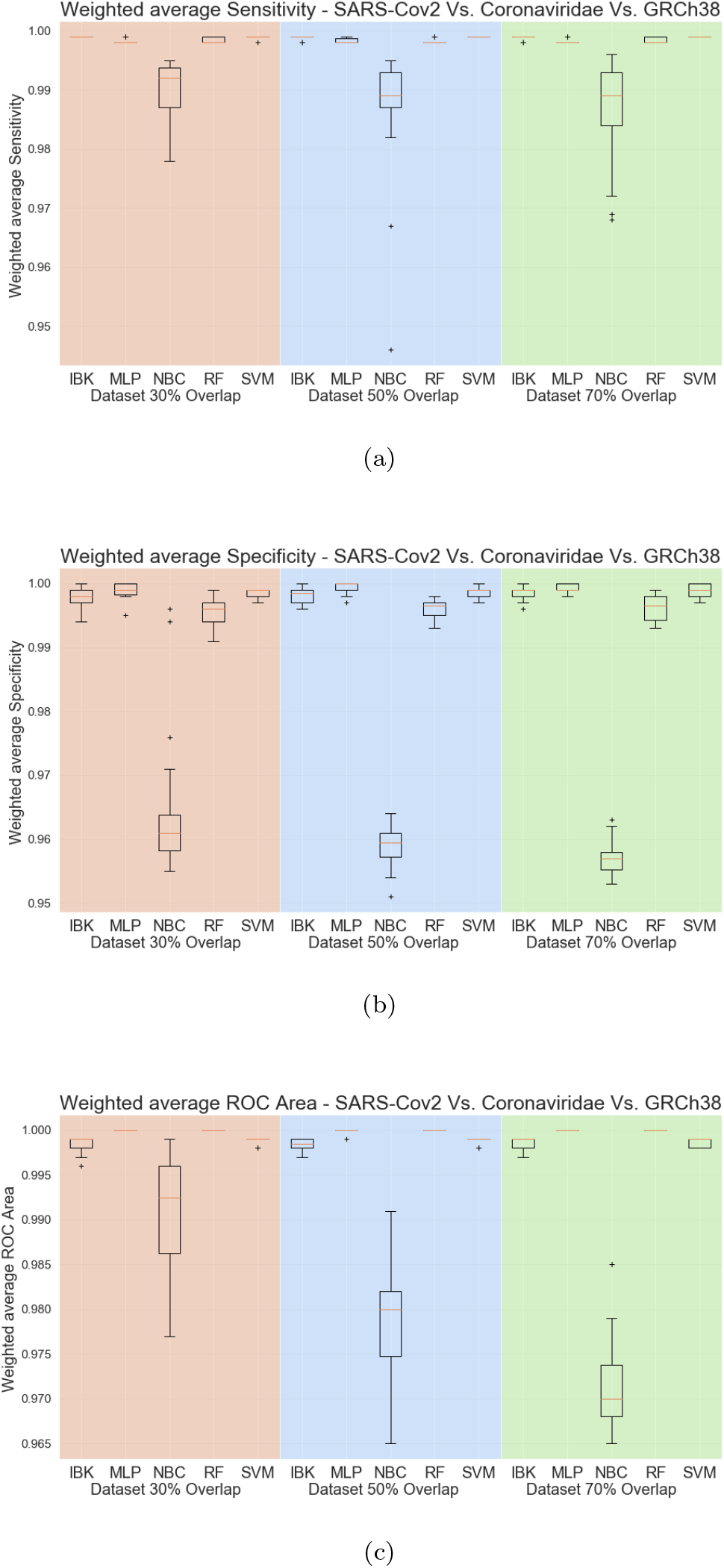
Weighted average sensitivity (a), specificity (b), and ROC area for SARS-Cov2 test scenario.

Aiming to better evaluate the results of the classifiers in the SARS-Cov2 test task, all the confusion matrices for IBK, MLP, Random Forest, and SVM classifiers are shown in Figure 21. IBK and Random Forest classifiers presents a confusion between SARS-Cov2 and Coronaviridae that varies from 10.26% (Figure 21-h) to 14.77% (Figure 21-c). This outcome is even worse for SVM classifier since most of the SARS-Cov2 examples are misclassified as Coronaviridae. By confusion matrix analysis, The MLP classifier has lower confusion rates between SARS-Cov2 and Coronaviridae. The results from MLP classifier in the dataset with 50% overlap (Figure 21-e) shows 99.92% average true positive rate for GRCh38 class, and 98.82% for the SARS-Cov2. For the Coronaviridae class, this classifier achieves 96.2%, while only 3.73% of Coronaviridae examples are misclassified as SARS-Cov2. Table 6 shows the sensitivity, specificity and ROC Area for each of the classes for this MLP classifier.

**Figure 21:**
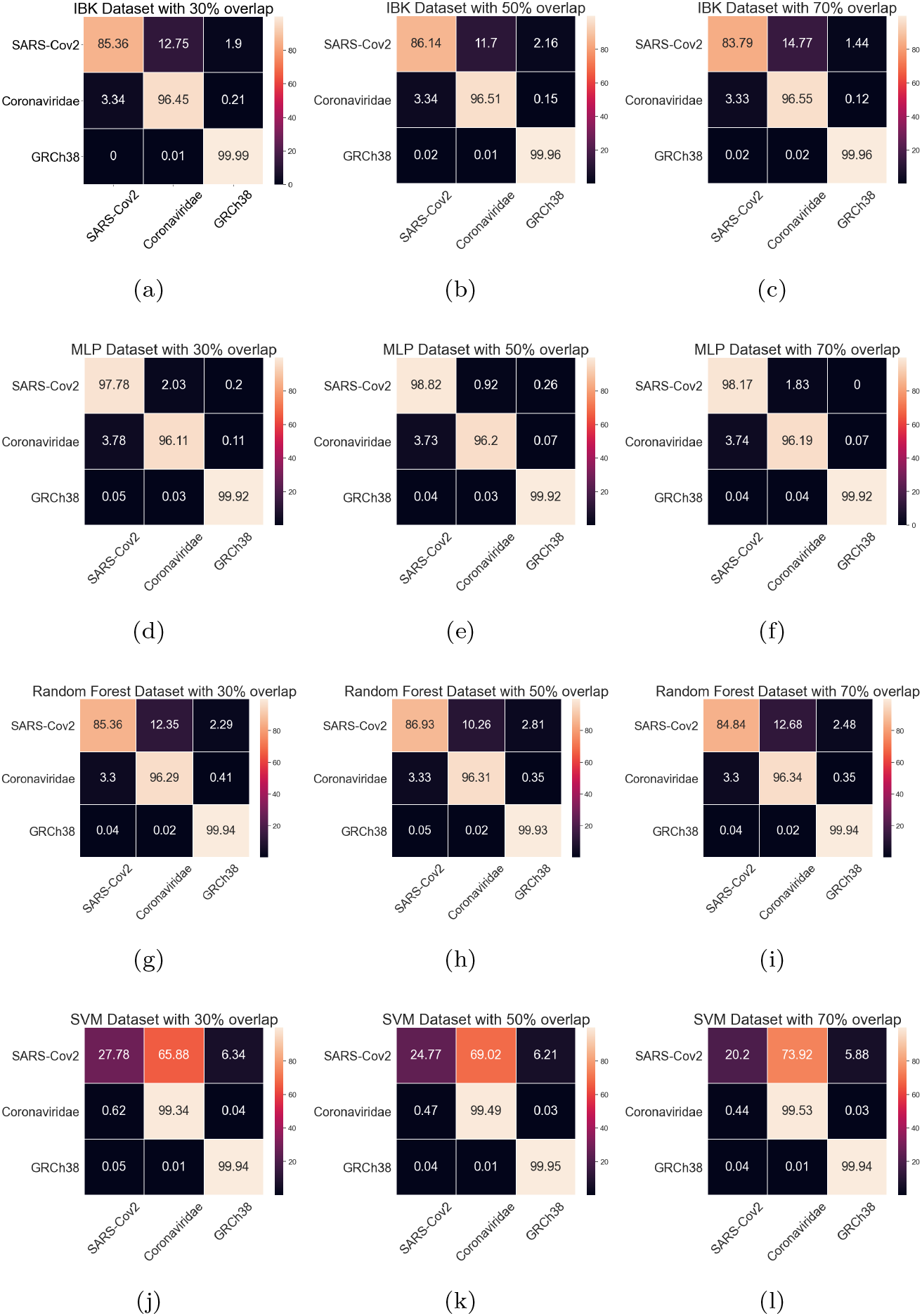
Average Confusion matrices for IBK, MLP, Random Forest and SMV in the SARS-Cov2 test scenario.

**Table 6:** Results of Sensitivity, specificity, and ROC area for MLP classifier in the SARS-Cov2 test scenario (results from dataset with 50% overlap).

| Class | Sensitivity-Recall |  | Specificity |  | ROC Area |  |
| --- | --- | --- | --- | --- | --- | --- |
|  | Average | Std. Dev. | Average | Std. Dev. | Average | Std. Dev. |
| **SARS-Cov2** | **0.98824** | **0.01198** | **0.99860** | **0.00020** | **0.99947** | **0.00056** |
| Coronaviridae | 0.96196 | 0.00190 | 0.99967 | 0.00017 | 0.99810 | 0.00094 |
| GRCh38 | 0.99923 | 0.00028 | 0.99928 | 0.00094 | 0.99997 | 0.00018 |

## 5. Discussion

Regarding the feature extraction methods, it seems to capture the structure of the viruses’ genome sequence. Random Forest classifier achieved the best overall performance for multiclass scenarios, while MLP classifier presented the best results for scenarios with fewer classes.

Evaluating the parameters for the feature extraction proposed method, splitting the viruses’ genome sequence into four folders (*n* = 4) seems to be enough to produce representative features. Regarding the overlap percentage, the proposed feature extraction method is not very sensitive to this parameter, even though 30% to 50% seems to be enough to produce good features representations.

The first multiclass scenario (with 25 viruses classes) is an extreme case scenario. Nevertheless, the Random Forest classifier achieved sensitivity and specificity above 0.9 for many classes. For those classes with lower sensitivity, the confusion matrix shows that most confusions are particular between two viruses. For example, Filoriviridae is the class with a lower sensitivity rate (0.23). However, checking the confusion matrix, on average, 76.27% of Filoriviridae are misclassified as Ebola Virus. There is no other significant confusion for Filoriviridae, so it is possible to design a classifier cascade to solve this specific confusion between two viruses.

One particular virus class is the Pharma Viridae since it has only 42 samples in the dataset (30 used for training and 12 for testing). Even with this small amount of samples in the multiclass scenario, the significant misclassifications for Pharma Viridae are Henteraviridae (22.78%), and Peribunyavirida (35.26%). With a larger sample size for the Pharma Viridae, classifiers could find a better boundary decision reducing this level of false-negative rate. However, for this particular class, three-classes cascade classifiers could be evaluated to deal with these types of errors.

Regardless of the feature extraction parameters or even the used classifier, there is still a 3-4% of Coronaviridae samples misclassified as SARS-Cov2. However, this is an expected outcome, since SARS-Cov2 belongs to the Coronaviridae family. Visualizing the extracted features, we found some samples of SARS-Cov2 and Coronaviridae that can not be distinguished, as showed in Figure 22. So, it is tough for any classifier to separate those two classes optimally.

**Figure 22:**
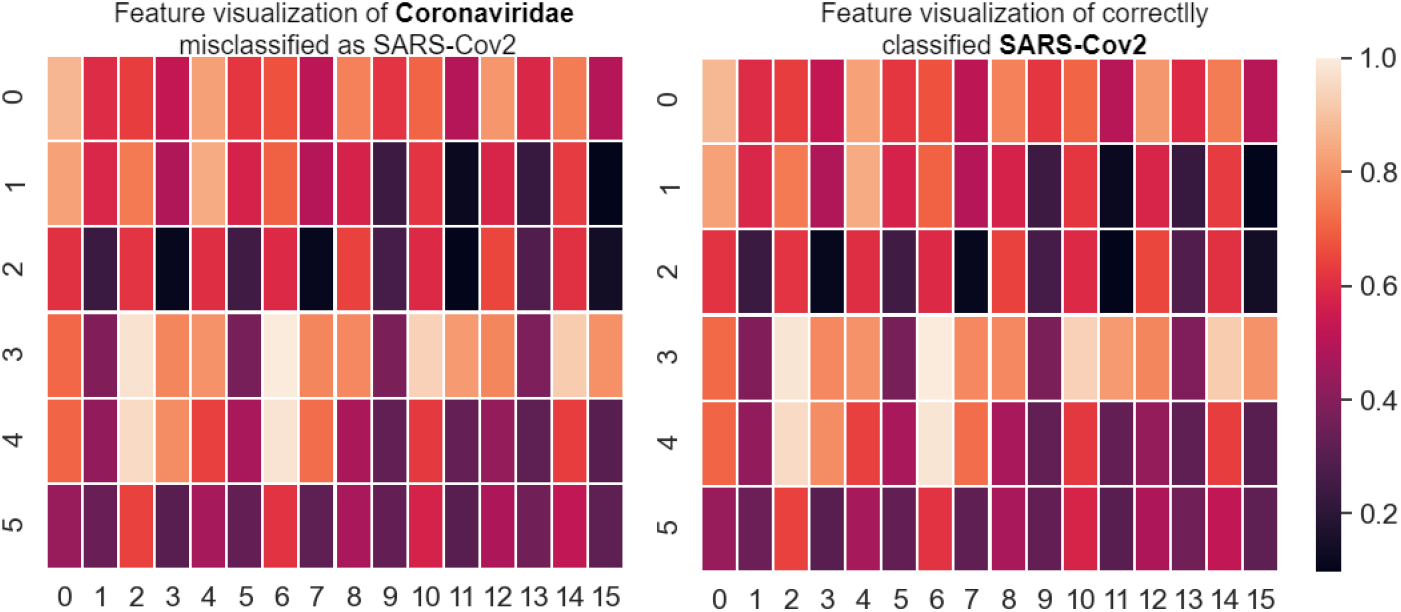
Feature visualization for selected SARS-Cov2 and Coronaviridae sample.

## 6. Conclusion

In this work we presented a novel method to represent DNA sequences by using pseudo-convolutions and co-occurrence matrices. With this method, we were able to represent hundreds of thousands of DNA sequences from 24 virus families. Then we separated SARS-Cov-2 sequences from the Coronaviridae family and demonstrated that our model is able to differentiate all virus families present on our database. SARS-Cov-2 was discriminated from virus families other than Coronaviridade and even from other coronaviruses with very high sensitivity and specificity.

We aimed to show the capabilities of optimizing the molecular diagnosis of Covid-19 by combining RT-PCR, the actual ground-truth Covid-19 diagnostic method, and our pseudo-convolutional method to identify SARS-Cov-2 DNA sequences faster.

From the obtained results, we can assume that the proposed pseudo-convolutional approach is able to characterize SARS-Cov-2 DNA sequences. This new representation of DNA sequences can be successfully used as a feature extraction stage to full connected networks, in order to use the deep learning philosophy, or other classical classification architectures. The evaluation of the proposed approach in real test scenarios, necessarily reduced to a limited set of virus families and healthy human sample DNA, showed high sensitivity (higher than 0.988) and specificity (higher than 0.998) rate as well. Hence, other researchers can use our solution and our methods to improve their results to diagnose Covid-19 faster with accuracies even higher than the state-of-the-art methods.

## Acknowledgements

We are grateful to the Brazilian research-funding agency CNPq, for the partial support of this research.

## Compliance with Ethical Standards

This study was partially funded by the Brazilian research agency Conselho Nacional de Desenvolvimento Científico e Tecnológico, CNPq.

All procedures performed in studies involving human participants were in accordance with the ethical standards of the institutional and/or national research committee and with the 1964 Helsinki declaration and its later amendments or comparable ethical standards.

